# Chromato-kinetic fingerprinting enables multiomic digital counting of single disease biomarker molecules

**DOI:** 10.1101/2025.01.31.636009

**Authors:** Pavel Banerjee, Sujay Ray, Liuhan Dai, Erin Sandford, Tanmay Chatterjee, Shankar Mandal, Javed Siddiqui, Muneesh Tewari, Nils G. Walter

## Abstract

Early and personalized intervention in complex diseases requires robust molecular diagnostics, yet the simultaneous detection of diverse biomarkers—microRNAs (miRNAs), mutant DNAs, and proteins—remains challenging due to low abundance and preprocessing incompatibilities. We present Biomarker Single-molecule Chromato-kinetic multi-Omics Profiling and Enumeration (Bio-SCOPE), a next-generation, triple-modality, multiplexed detection platform that integrates both chromatic and kinetic fingerprinting for molecular profiling through digital encoding. Bio-SCOPE achieves femtomolar sensitivity, single-base mismatch specificity, and minimal matrix interference, enabling precise, parallel quantification of up to six biomarkers in a single sample with single-molecule resolution. We demonstrate its versatility in accurately detecting low-abundance miRNA signatures from human tissues, identifying upregulated miRNAs in the plasma of prostate cancer patients, and measuring elevated interleukin-6 (IL-6) and hsa-miR-21 levels in cytokine release syndrome patients. By seamlessly integrating multiomic biomarker panels on a unified, high-precision platform, Bio-SCOPE provides a transformative tool for molecular diagnostics and precision medicine.

## Introduction

The advent of advanced molecular diagnostics has transformed biomarker detection, providing critical insights into disease mechanisms and enabling early diagnosis and personalized treatments. Key biomarkers—such as microRNAs (miRNAs), DNA mutations, and proteins—play a crucial role in managing complex conditions, including cancer^1^, cardiovascular diseases^2^, and neurodegenerative disorders^3^. Biological and pathological processes often influence the levels of multiple biomarkers at once, making technologies for multiplexed detection particularly valuable for the increased sensitivity and/or specificity they provide^4,5^. For example, liquid biopsies, which analyze molecular biomarkers of cancer in biofluids like serum and plasma, enable the simultaneous measurement of multiple analytes from a single sample, offering a comprehensive molecular portrait with enhanced diagnostic and prognostic value^6–9^.

Despite these advancements, practical implementation of multiplexed assays faces significant challenges. Assays of biological samples frequently exhibit cross-reactivity and nonspecific binding of probes, resulting in ambiguous or misleading results^10^. Detecting low-abundance targets amidst prevalent molecules necessitates sophisticated analytical techniques and robust signal amplification^11^. Established nucleic acid techniques such as polymerase chain reaction (PCR) often suffer from amplification bias^11,12^, while next-generation sequencing (NGS) and nanopore sequencing face challenges with sequence specificity and sensitivity^13,14^. Amplification free nucleic acid assays^15–17^ exhibit high false-positive rates and constraints due to degenerate hybridization thermodynamics^18,19^, limiting them to picomolar sensitivity and thus only able to detect high-abundance targets. Additionally, these methods are not applicable to other analyte classes such as proteins^11^. Innovative barcoding methods, such as DNA origami^20^ or molecular indexing^21^, require extensive data analysis and bioinformatics expertise, posing barriers to clinical integration. These challenges in balancing sensitivity, specificity, cost, speed, and scalability underscore the need for further innovation in multiplexed and multiomic detection for clinical applications.

Single-molecule fluorescence offers high sensitivity without amplification, potentially offering a universal multiomic platform. However, multiplexing faces challenges such as incomplete control over labeling and bleaching, as well as spectral overlap, which limits the number of colors that can be distinguished^22,23^. Methods like Exchange-PAINT and MERFISH overcome this problem by using sequential probe exchange, albeit at the cost of greater acquisition time^24–26^. FRETfluors expand color-based multiplexing through the use of ratiometric spectroscopy and fluorescence lifetime^9^. However, for all these methods using conventional tightly binding probes, specificity is still limited by binding thermodynamics, making it challenging to detect single-nucleotide mutations with high specificity^18^.

To overcome these limitations, technologies for single-molecule fluorescent multiplexing offer unique “fingerprints” for analytes that enable high-specificity detection with minimal interference exploiting temporal^26^, spatial^15,20^ and kinetic dimensions^22,24,27–30^. Kinetic fingerprinting leverages binding dynamics of biomolecular interactions, distinguishing targets based on kinetic profiles^28^. It offers ddPCR-like specificity^29^ as well as single molecule sensitivity for nucleic acid^28^ and proteins^30,31^, although multiplexing has not been demonstrated for these assays. Color-based fingerprinting employs distinct signals for analytes, simplifying multiplexed detection but facing limitations similar to fluorescence-based techniques, and practically is limited to only ∼3 distinguishable signatures on experimentally accessible timescales^22^.

Here, we present Biomarker Single-molecule Chromato-kinetic multi-Omics Profiling and Enumeration (Bio-SCOPE), a multiplexed analyte detection strategy that leverages both chromatic and kinetic fingerprint encoding to achieve high-specificity, amplification-free detection of diverse analytes. Our method achieves femtomolar sensitivity and single-base mismatch selectivity, while enabling―since it does not rely on PCR amplification―the simultaneous analysis of multiple classes of disease biomarkers, including miRNAs, mutant DNAs, and proteins. We demonstrate the digital single-molecule detection and quantification of up to six targets simultaneously from a single sample, the detection of low-abundance targets amidst high-abundance targets, and the quantitative mapping of multiple miRNAs across diverse human tissues. Using Bio-SCOPE, we observe the specific upregulation of miRNAs *hsa*-miR-141 and *hsa*-miR-375 in prostate cancer patient plasma samples. Further, we detect the selective upregulation of interleukin-6 (IL-6) and *hsa*-miR-21, but not interleukin-8 (IL-8) or *hsa*-miR-16, in the serum of patient with cytokine release syndrome (CRS). These findings indicate that Bio-SCOPE, unlike existing approaches, generates simultaneous, internally calibrated, high-sensitivity multiomic biomarker profiles of patients, which could be vital for early disease detection, progression monitoring, and treatment assessment, thereby advancing clinical diagnostic workflows with potential for revolutionizing precision medicine through deeper insights for clinicians.

## Results

### Overall strategy and optimization of Bio-SCOPE fingerprinting

The Bio-SCOPE workflow encompasses biofluid sample collection and the determination of target abundance through the recognition of unique kinetic fingerprints of transient, repeated interactions between probe and analyte. As a first proof-of-concept, we aimed to assay human serum, plasma, and tissue-derived RNA extracts by Total Internal Reflection Fluorescence (TIRF) microscopy to capture these kinetic fingerprints—temporal binding patterns—that sensitively distinguish specific target interactions from nonspecific interactions with assay surfaces or other matrix components (Fig. 1a(i,ii); Extended Data Fig. 1). This differentiation is achieved by applying specific filtering parameters such as minimum and/or maximum thresholds with respect to the number of binding and dissociation events (N_b+d_), median dwell time in the bound state (τ_on,med_) and median dwell time in the unbound state (τ_off,med_; Supplementary Table 2). Without kinetic filtering―not possible for thermodynamic binding-driven single molecule detection approaches―nonspecific binding predominates among the single-molecule traces, causing numerous false positives per field of view (FOV; Extended Data Fig. 1d). Kinetic filtering reduces these false positives to nearly zero (<1 per FOV) while preserving most true positives, underscoring the high specificity of single-molecule detection via kinetic fingerprinting. For multiplexing, the methodology leverages three fluorescence channels—Cy3 (cyan, C_1_), Cy5 (magenta, C_2_), and mixed Cy3/Cy5 binding (dark yellow, C_12_)—to combine chromatic fingerprints with kinetic behaviors (Fig. 1a(iii), Fig. 1a(iv)). This approach is expected to allow the concurrent quantification of multiple targets in complex biofluid mixtures, reflecting real-world diagnostic requirements (Fig. 1a(v)).

**Fig. 1.**
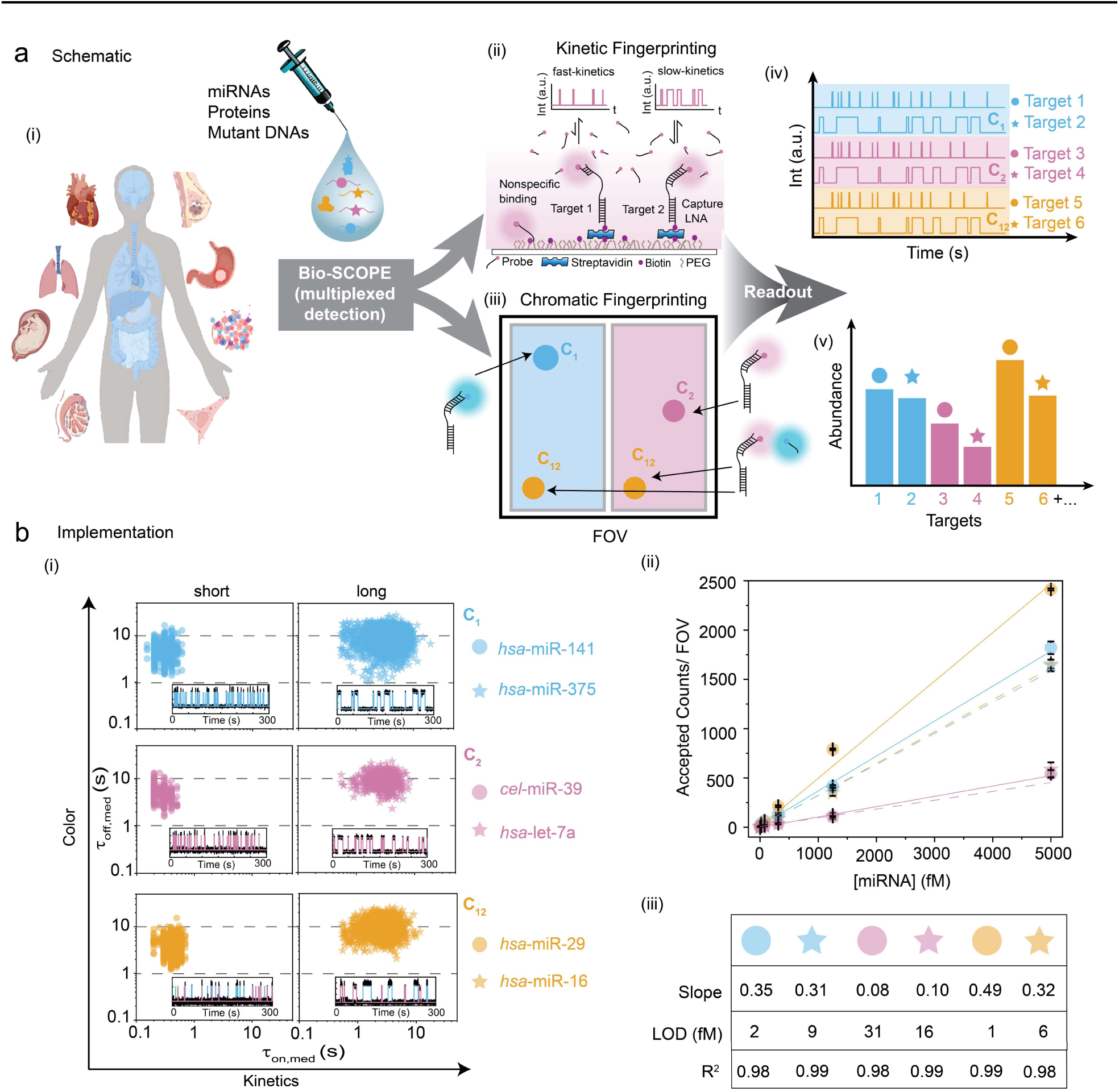
ǀ Principle of Bio-SCOPE for the quantitative detection of diverse targets. **a**, Overall workflow for multiplexed detection and quantification of key biomarkers, such as miRNAs, mutant DNAs, and proteins in biofluids by Bio-SCOPE (i) Samples are collected from various types of human biofluids relevant to liquid biopsies, such as serum, plasma, and RNA extracts from different human tissues and organs. (ii) Different interaction of a kinetic fingerprinting probe with a surface-captured target yields diverse temporal patterns of repeated binding and dissociation, or kinetic fingerprints, distinct from nonspecific interactions of probes with surface or matrix contaminants. These kinetic fingerprints are measured by TIRF microscopy. (iii) Defining three color channels (C_1_= only Cy3 (cyan), C_2_= only Cy5 (magenta) and C_12_= combination of Cy3 and Cy5 (dark yellow)) of diverse temporal patterns of repeated binding and dissociation, or chromatic fingerprinting. (iv) Schematic view of chromato-kinetic fingerprints indicative of repetitive fluorescent probe binding in different color channels and with distinguishable kinetics. (v) Schematic of relative abundance assessments of targets by Bio-SCOPE. **b,** (i) Experimental assay optimization and channelization of six synthetic miRNAs to identify shared preparatory and imaging conditions. Scatterplots of dwell time analysis (i.e., τ_on,med_ versus τ_off,med_) for all accepted intensity-versus-time trajectories observed in a single FOV for each of the six exemplary miRNAs (*hsa*-miR-141, hsa-miR-375, *cel*-miR-39, *hsa*-let-7a, *hsa*-miR-29, *hsa*-miR-16), optimized separately. Representative single molecule trace behaviors are shown in the insets. Target concentrations: 5 pM; Temperature: 25 °C; Acquisition time: 5 min/FOV; Probes: *hsa*-miR-141_NS_DNA_FP_Cy3_8-nt at 50 nM, hsa-miR-375_NS_DNA_FP_Cy3_C8-nt at 50 nM, *cel*-miR-39_NS_DNA_FP_Cy5_8-nt at 50 nM, *hsa*-let-7a_NS_DNA_FP_Cy5_11-nt at 50 nM, *hsa*-miR-29_NS_DNA_FP_Cy3 & Cy5_8-nt at 25 nM (each), *hsa*-miR-16_NS_DNA_FP_Cy3 & Cy5_C10-nt at 25 nM (each). (ii) Standard curves for the six miRNAs after optimization using separate fingerprinting assays. Linear regression fits are shown as solid or dashed lines. Error bars indicate the standard deviation (SD) of three independent measurements. (iii) Slopes from standard curves, estimated LOD values (in fM) and R^2^ values from fitting for the six miRNAs.

Details of the experimental implementation of Bio-SCOPE, assay optimization and analysis pipeline are found in ‘Methods’. Probes against five synthetic human miRNAs that are known to be dysregulated biomarkers in various cancers^32,33^ (*hsa*-miR-141, *hsa*-miR-375, *hsa*-let-7a, *hsa*-miR-29, and *hsa*-miR-16), and one internal control miRNA from *C. elegans* (*cel*-miR-39)^4^, were individually optimized for assaying under standardized conditions to produce distinct chromato-kinetic signatures, evidenced by distinguishable scatterplots of median dwell times (τ_on,med_ versus τ_off,med_) derived from single-molecule traces (Fig. 1b(i)). Standard curves produced from these individual assays exhibit strong linearity and reproducibility (Fig. 1b(ii)), validating the quantitative robustness of Bio-SCOPE for potential use in multiplexed applications. An exponential cumulative distribution function (CDF), fitted to dwell times of thousands of probe-bound and probe-unbound events per condition from over a thousand molecules, precisely estimates ensemble-averaged <τ_on_> and <τ_off_> values, respectively (Extended Data Fig. 1e-g). The assays exhibit high sensitivity, with low-femtomolar limits of detection (LODs; Fig. 1b(iii)).

Distinct kinetic fingerprints suitable for multiplexing can be achieved by tuning a range of experimental variables, such as the sequence, length, and concentration of fingerprinting probes (FPs), measurement temperature, fluorescence dye used, and chemical makeup of the FPs (DNA, RNA, with or without chemical modifications; Extended Data Fig. 2). These different assay conditions yield distinct <τ_on_> and <τ_off_> values from single-exponential fittings of CDFs (Extended Data Fig. 2h), illustrating several degrees of freedom for fine-tuning kinetic assay parameters to achieve highest performance in Bio-SCOPE.

### Kinetic and chromatic multiplexing each distinguishes synthetic miRNAs in a FOV

Next, we investigated Bio-SCOPE’s capacity for multiplexing in the kinetic dimension. The classification of dwell times for multiplexed datasets from a single FOV (Fig. 2a) can be achieved using *k*-means clustering, with the clustering model pretrained from experiments with the individual targets whose kinetics are designed to be distinguishable in a single-color channel (Fig. 2b). Simulations demonstrated a significant reduction in unassigned molecules with increasing acquisition time, reaching a point of diminishing improvement around 300 s, guiding the choice of optimal experimental times and clustering parameters (Extended Data Fig. 3a). A robustness of 95% confidence in clustering was further validated for simulated traces through the multiplexing of three targets at extreme acquisition times (50 s and 1,000 s, Extended Data Fig. 3b).

**Fig. 2.**
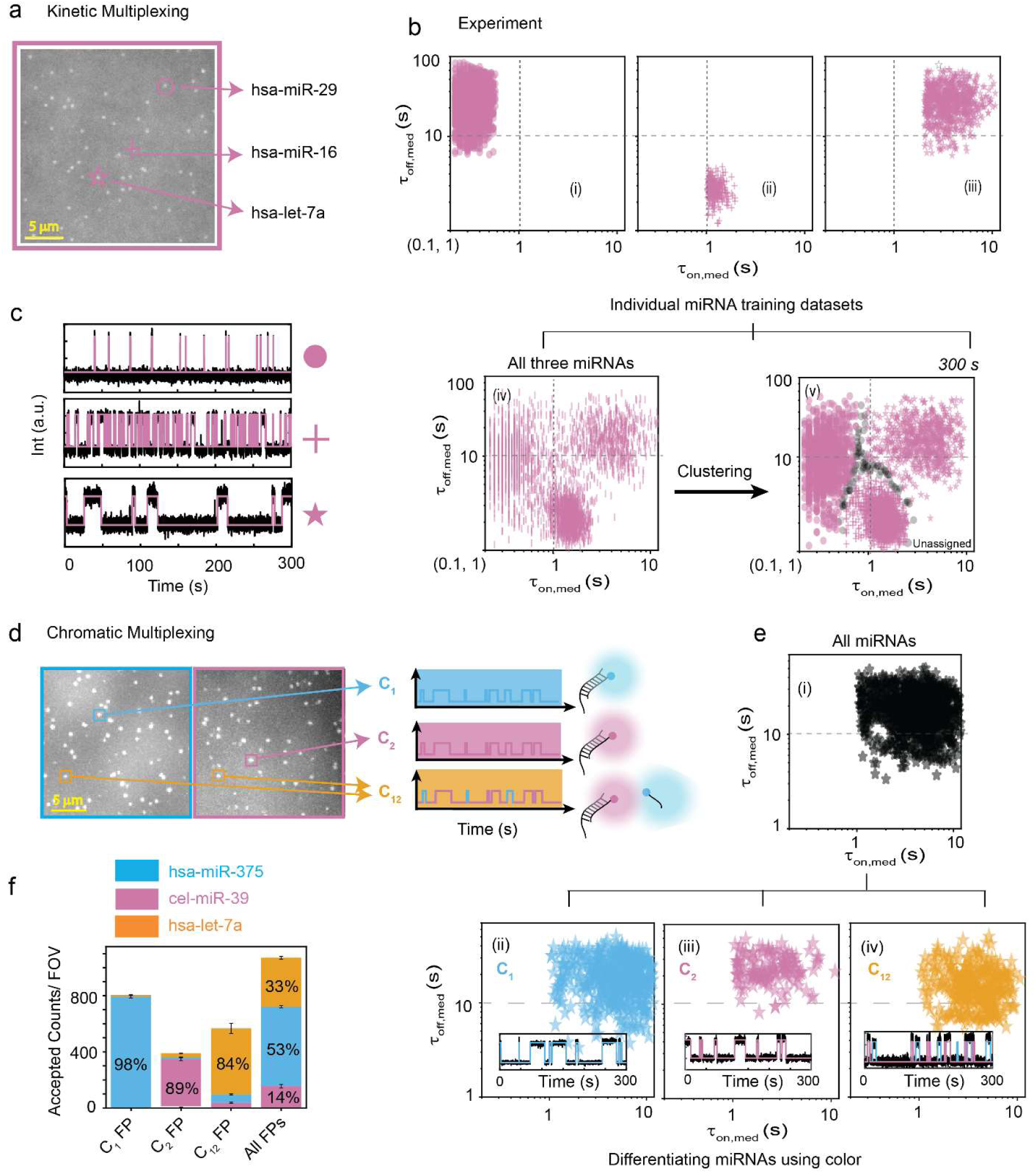
ǀ Demonstration of kinetic and chromatic multiplexing. **a**, Average movie of a representative portion of a TIRF microscope FOV (scale bar: 5 µm), showing bright puncta at the locations where single fluorescent probes are bound at or near the imaging surface and representative kinetic fingerprinting spots of three defined kinetics. **b,** Scatterplots of dwell time analysis (i.e., τ_on,med_ versus τ_off,med_) for all accepted time trajectories observed within five FOVs of (i) *hsa*-miR-29, (ii) *hsa*-miR-16, (iii) *hsa*-let-7a, (iv) all three miRNAs in a multiplex assay (without clustering), and (v) the same with 95% confidence clusters (N_unassigned_ = 2%). Unassigned molecules during the clustering are represented by black circles. **c,** Representative kinetic fingerprinting (i.e., single-molecule intensity-versus-time traces from TIRF microscopy measurements) of (i) *hsa*-miR-29 (short τ_on_, long τ_off_), (ii) *hsa*-miR-16 (intermediate τ_on,_ short τ_off_), (iii) *hsa*-let-7a (long τ_on_, long τ_off_). Raw intensity-versus-time traces (black lines) were idealized by hidden Markov modeling (magenta lines) to extract kinetic parameters for analysis. Target concentration ratio: 1:1:5; Probes: *hsa*-miR-29_NS_DNA FP_Cy5_8-nt at 25 nM, *hsa*-miR-16_NS_DNA_FP_Cy5_C9-nt at 75 nM, *hsa*-let-7a_NS_DNA_ FP_Cy5_11-nt at 25 nM. **d,** Average movie of a representative portion of a TIRF microscope FOV (scale bar: 5 µm), divided into two channels, showing bright puncta at the locations where single fluorescent probes are bound at or near the imaging surface and representative schematic chromatic fingerprinting of three defined color channels. **e,** Demultiplexing of three miRNAs using color: (i) all three miRNAs in multiplexed condition with similar τ_on,med_ and τ_off,med_ and, demultiplexed (ii) *hsa*-miR-375 (C_1_ channel), (iii) *hsa*-cel-39 (C_2_ channel), and (iv) *hsa*-let-7a (C_12_ channel). Representative single molecule trace behaviors are shown in the insets. **f,** Relative abundance of three miRNAs in sequential and all-in-one sample multiplexing condition using chromatic fingerprinting. Error bars indicate the SD of three independent measurements. Target concentration ratio: 1:5:5; Probes: *hsa*-miR-375_NS_DNA_ FP_Cy3_C8-nt at 30 nM, *cel*-miR-39_NS_DNA_FP_Cy5_10-nt at 30 nM, *hsa*-let-7a_NS_DNA_ FP_Cy5 & Cy3_11-nt at 30 nM.

We validated our clustering approach for distinguishing experimental target signals using scatterplots of median FP dwell times for individual synthetic targets (*hsa*-miR-29, *hsa*-miR-16, and *hsa*-let-7a) as well as a mixture of all three measured simultaneously (Fig. 2b(i–v)). Reflecting distinct temporal patterns of FP binding, *hsa*-miR-29 detection exhibited the shortest ensemble <τ_on_> (0.49 ± 0.02s; i.e., fastest dissociation) and a long <τ_off_> (10.35 ± 0.07s), *hsa*-miR-16 displayed an intermediate <τ_on_> (2.29 ± 0.04s) combined with the shortest <τ_off_> (2.69 ± 0.01s) due to its chosen high FP concentration, and *hsa*-let-7a showed the longest <τ_on_> (6.25 ± 0.05s; i.e., slowest dissociation) with the longest <τ_off_> (16.47 ± 0.05s) (Fig. 2c, Supplementary Fig. 1). The robustness of Bio-SCOPE for multiplexed, simultaneous kinetic detection was further evidenced by clustering at different confidence levels (90%, 95%, and 99%) when two miRNAs were mixed (Extended Data Fig. 4). Notably, kinetic multiplexing at a 95% confidence level effectively resolved all pairwise mixtures of the three miRNAs at the single target molecule level within the same FOV (Supplementary Fig. 1c).

We similarly tested chromatic multiplexing using our three-color channels. Based on the signals from Cy3, Cy5, or a mixture of both dyes in a given location within a FOV, a detected molecule could be assigned to one of these three channels (C_1_ = *hsa*-miR-375, C_2_ = *cel*-miR-39, C_12_ = *hsa*-let-7a; Fig. 2d, Supplementary Fig. 2), despite overlapping τ_on,med_ and τ_off,med_ distributions (Fig. 2e, Supplementary Fig. 3). The observed consistent relative abundances of these miRNAs when measured separately or in an equimolar mixture supports the robustness of simultaneous chromatic fingerprinting (Fig. 2f). Combining kinetic and chromatic discriminators thus enables the parallel, high-confidence quantification of up to nine biomarkers in a single sample with single-molecule sensitivity.

### Bio-SCOPE achieves robust chromato-kinetic multiplexing with femtomolar sensitivity

To fully realize Bio-SCOPE and its multiplexing capabilities, we integrated kinetic and chromatic multiplexing and performed initial testing on synthetic mixtures of all six miRNAs. 3D scatterplots of τ_on,med_, τ_off,med_ and color simultaneously resolved the six targets within the same FOV into distinct clusters of molecules at 95% confidence, with 4-8% of molecules remaining unassigned (Fig. 3a; Supplementary Fig. 4).

**Fig. 3.**
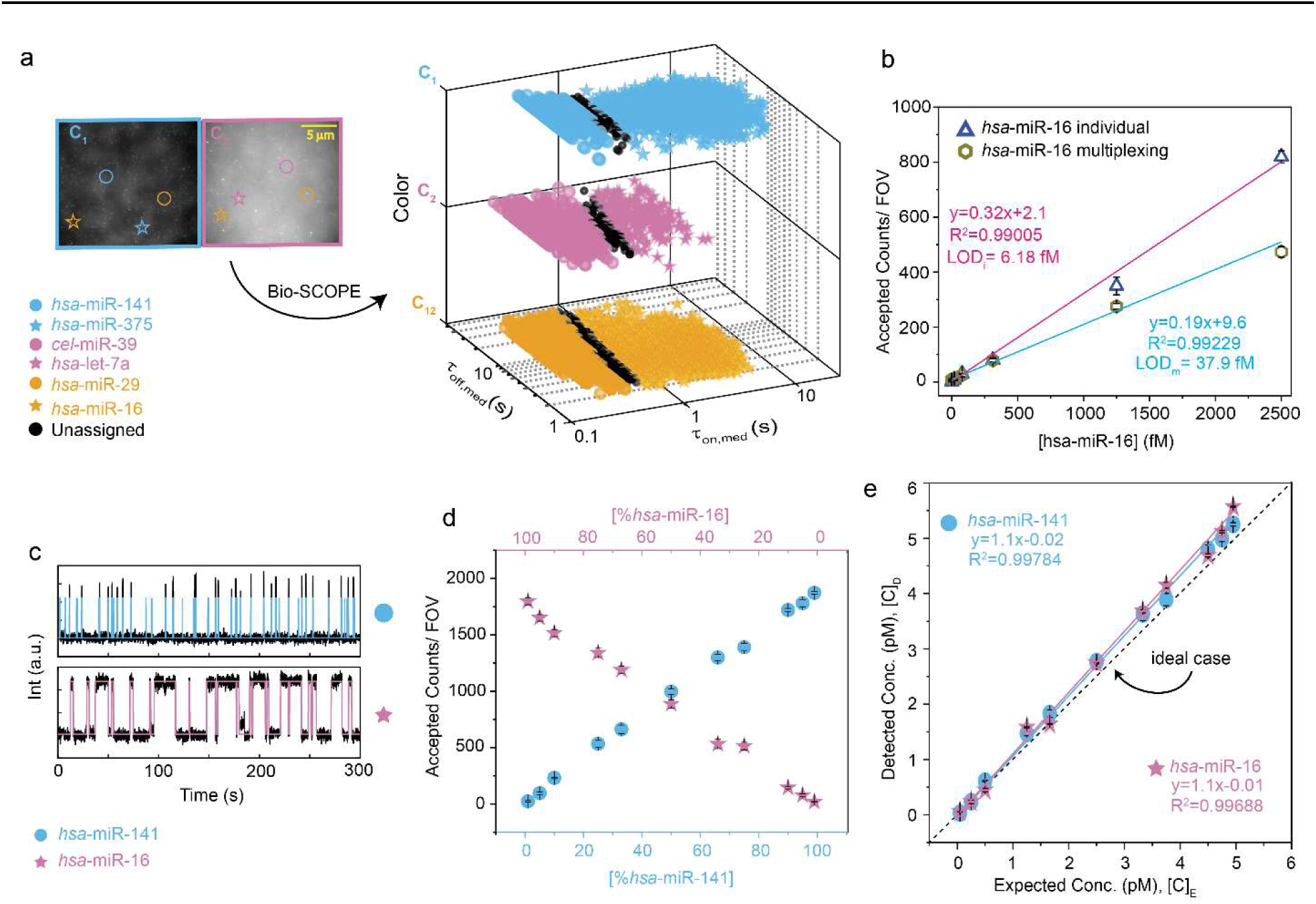
ǀ Multiplexed detection and quantification of miRNAs by Bio-SCOPE. **a**, Average movie of a representative portion of a TIRF microscope FOV (scale bar: 5 µm), divided into two channels, showing chromato-kinetic fingerprinting using Bio-SCOPE and 3D scatterplot of dwell time analysis (i.e., τ_on,med_ versus τ_off,med_ with color channels) for all accepted intensity-versus-time trajectories observed within five FOV (with 95% confidence clusters). Unassigned molecules during the clustering are represented by black circles. Target concentration ratio: 1:1:5:5:1:1; Probes: *hsa*-miR-141_NS_DNA_FP_Cy3_8-nt at 25 nM, *hsa*-miR-375_NS_DNA_FP_Cy3_C8-nt at 75 nM, *cel*-miR-39_NS_DNA_FP_Cy5_8-nt at 25 nM, *hsa*-let-7a_NS_DNA_FP_Cy5_11-nt at 75 nM, *hsa*-miR-29_NS_DNA_FP_Cy3 & Cy5_8-nt at 12.5 nM (each), *hsa*-miR-16_NS_DNA_FP_Cy3 & Cy5_C10-nt at 37.5 nM (each). **b,** Standard curves of *hsa*-miR-16 in individual and multiplexing scenarios. Linear regression fits are shown as solid lines. Fitting equations, R^2^ values and estimated LODs are documented within the figure. **c,** Representative single molecule trace behaviors of the detection of two similarly abundant miRNAs (i.e., *hsa*-miR-141 and *hsa*-miR-16) using Bio-SCOPE, where *hsa*-miR-141 is placed in the C_1_ channel with short τ_on_ and *hsa*-miR-16 is in the C_2_ channel with long τ_on._ **d,** Detection of *hsa*-miR-141 and *hsa*-miR-16 in different ratios using Bio-SCOPE. Total target concentration: 5 pM. **e,** Expected concentration versus detected concentration plot of the ratio-metric datasets from (d). Detected concentrations are estimated from the accepted counts obtained per FOV (normalized by the slope of the standard curves of individual miRNAs). Linear regression fits are shown as solid lines. The dashed line indicates the ideal scenario of detecting exact concentration as expected concentration. Black error bars represent the standard errors of the mean from three independent replicates.

To test the impact of multiplexing on sensitivity, we compared the LODs for the detection of targets individually and in the parallel 6-plex measurement. Standard curves for *hsa*-miR-16 alone and in the presence of a constant concentration of the five other miRNAs revealed a six-fold increase in LOD in the multiplexed measurement (though still in the low fM range; Fig. 3b). This increase is consistent with the expected six-fold drop in the surface density of each capture probe in the multiplexed measurement compared to the individual assays. As expected for a robust assay, the five other miRNAs were detected at constant concentrations (CV = 4-10%) in the presence of increasing *hsa*-miR-16 concentrations (Extended Data Fig. 5a). These results support the potential of Bio-SCOPE for sensitive, multiplexed biomarker quantification.

Next, we evaluated the accuracy of Bio-SCOPE in quantifying two miRNAs at varying ratios, as might be expected for patient samples. *hsa*-miR-141 and *hsa*-miR-16 were mixed in ratios ranging from 1:99 to 99:1 while maintaining a total miRNA concentration of 5 pM, then quantified by Bio-SCOPE. Representative single-molecule traces showed *hsa*-miR-141 in the C_1_ channel (short <τ_on_>= 0.26 ± 0.01s) and *hsa*-miR-16 in the C_2_ channel (long <τ_on_>= 2.95 ± 0.02s; Fig. 3c), each exhibiting distinctive kinetics of probe binding and dissociation (Extended Data Fig. 5b-c). Copies of each miRNA were detected in proportion to its prevalence in the mixture (Fig. 3d), yielding high concordance between the expected and measured concentrations of both miRNA targets, underscoring the method’s quantitative accuracy (Fig. 3e). Additionally, we demonstrated our ability to tune an assay to compensate for different abundances of miRNA targets by modifying the ratio of the respective capture probes on the surface. This approach allowed for optimized simultaneous quantification of, e.g., high-abundance *hsa*-miR-29 and low-abundance *hsa*-let-7a in a sample (Extended Data Fig. 5d), further demonstrating the overall robustness and adaptability of Bio-SCOPE.

### Bio-SCOPE achieves high single-nucleotide variant specificity in total cellular RNA with femtomolar sensitivity

An estimated ∼2,600 human miRNAs exert major post-transcriptional gene regulatory effects under both physiological and pathological conditions^34^, forming RNA-induced silencing complexes (RISC) with their effector protein Ago2 to mediate sequence-specific RNA silencing^33^. These miRNAs group into ∼260 families, distinguished by their conserved seed sequences, nucleotides 2-8 serving as core element for binding a regulated mRNA^35^. Members of a miRNA family exhibit extremely high sequence homology, often differing by just one nucleotide outside the seed sequence^7,28^, making it challenging to distinguish them. To demonstrate Bio-SCOPE’s specificity for single-nucleotide variants (SNVs), we therefore focused on multiplexed detection of three *hsa*-let-7 miRNAs family members of relatively high and similar abundance in human cells―let-7a, let-7b and let-7d―differing by either one or three nucleotides (Fig. 4a). As the ultimate challenge, a single probe was designed to interact differentially with these miRNAs (Fig. 4a), to be read out as kinetic patterns to be resolved using *k-*means clustering (Fig. 4b). Using a single probe reduced the opportunity to vary τ_off_ between clusters, necessitating clustering at a lower confidence level of 85% compared to our previous measurements (Supplementary Fig. 5). As predicted, representative kinetic fingerprints showed distinct patterns of repetitive probe binding to *hsa*-let-7b (short <τ_on_> = 0.10 ± 0.01s), *hsa*-let-7a (intermediate <τ_on_>= 0.86 ± 0.01s), and *hsa*-let-7d (long <τ_on_> = 11.34 ± 0.07s; Fig. 4c).

**Fig. 4.**
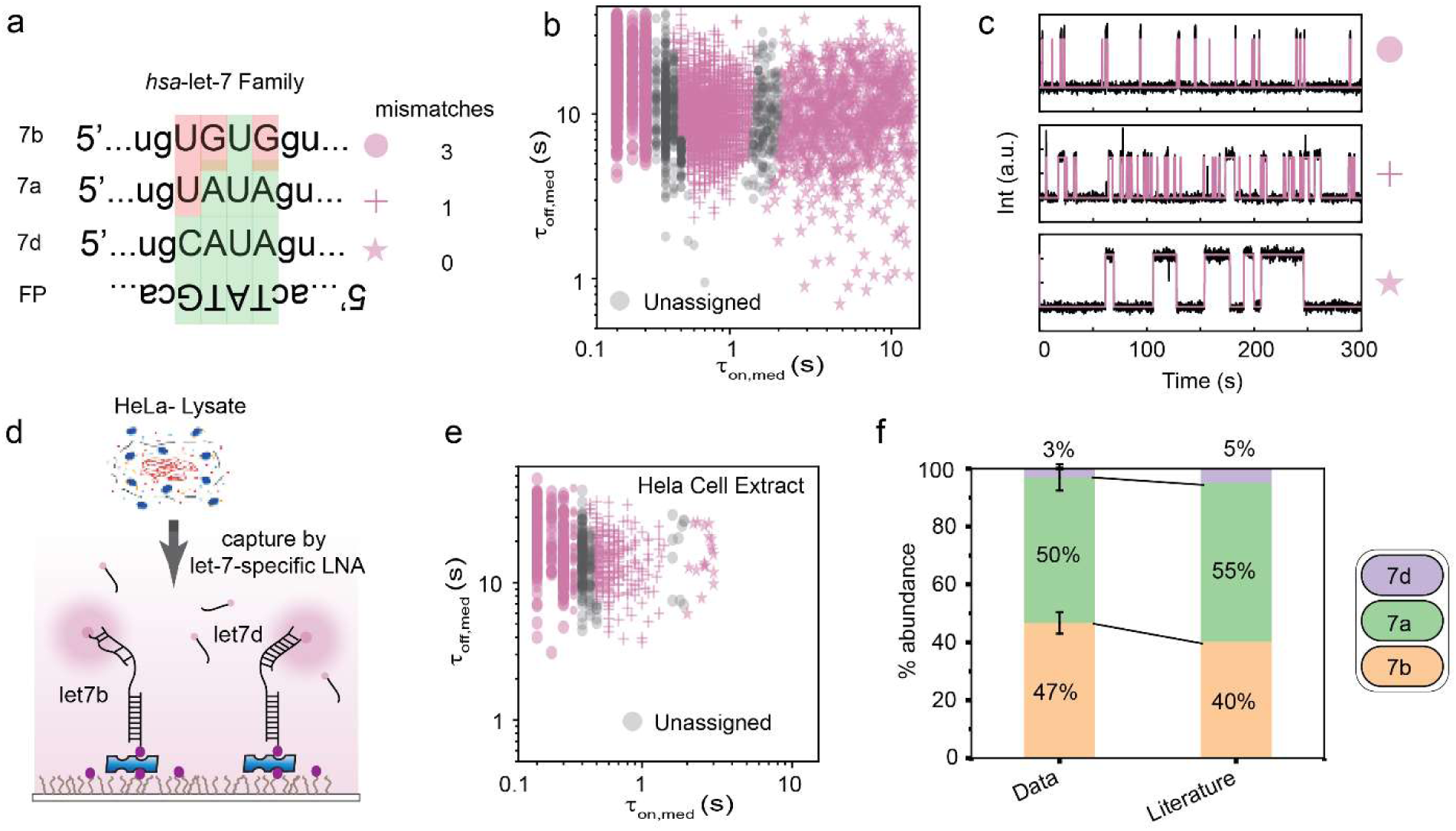
ǀ Multiplexed detection of miRNA family members representing single-nucleotide variants (SNVs). **a**, Sequence information of different let-7 family members (particularly in the single nucleotide variant region) and the designed common detection probe (FP). The number of mismatches between the probe and each target is listed. **b,** Kinetic multiplexing of the let-7 family three-plex dataset with 85% confidence clusters (5 FOVs, N_unassigned_ =13%). Unassigned molecules during the clustering are represented by black circles. **c,** Representative kinetic fingerprints indicative of repetitive probe binding (same fluorophore) to the different let-7 family members i.e., *hsa*-let-7b (short τ_on_), *hsa*-let-7a (intermediate τ_on_), and *hsa*-let-7d (long τ_on_). **d,** Schematic representation of kinetic multiplexing of the let-7 family in Hela cell extract using different number of mismatches. **e,** Kinetic multiplexing of the let-7 family three-plex dataset in Hela cell extract with 85% confidence clusters (5 FOVs, N_unassigned_ =14%). **f,** Relative abundance of three let-7 family miRNAs in Hela cell extract determined using kinetic multiplexing and comparison with previous reports^36^. Error bars indicate the SD of three independent measurements. Total target concentration: 12 pM (1:1:1); Probe: *hsa*-let-7_NS_DNA_FP_Cy5_11-nt at 50 nM.

To test its ability to quantify endogenous analytes within a complex mixture, we extended the Bio-SCOPE let-7a/b/d assay to a HeLa total RNA cell extract. By employing specific probe region blockers against other let-7 family members, we further enhanced target miRNA specificity (Fig. 4d). All three target miRNAs were detected by kinetic multiplexing (Fig. 4e), with relative abundances closely matching those reported in the literature (Fig. 4f)^36^. These results illustrate that Bio-SCOPE enables the accurate multiplexed quantification of closely related miRNAs, even against a background of thousands of other sequences in a total RNA preparation.

### Bio-SCOPE multiplexing accurately detects miRNA profiles in tissue and patient samples

To evaluate Bio-SCOPE’s potential for multiplexed biomarker detection for disease diagnostics, we profiled multiple miRNAs in total RNA extracts from diverse human tissue samples and cultured cell lines. We detected, and compared the relative expression levels of, our six miRNAs (*hsa*-miR-141, *hsa*-miR-375, *cel*-miR-39, *hsa*-let-7a, *hsa*-miR-29, and *hsa*-miR-16) across eight different human samples, including four healthy tissues (heart, prostate, lung, placenta) and four cancer cell lines (K562, HL-60, Raji, HeLa-S3; Fig. 5a). To ensure accurate quantification, measured miRNA concentrations were normalized against *C. elegans cel*-miR-39 (Supplementary Fig. 6)^4^; since *cel*-miR-39 is orthogonal to any human miRNAs, we doped *cel*-miR-39 at a constant, known concentration into each tissue sample, mitigating experimental variations caused by varying analyte matrices, as also done for PCR-based amplification assays^4^. The heat map of normalized concentrations [C]_N_ revealed distinct miRNA expression profiles across different tissue types (Fig. 5a). Comparative analysis highlighted significant differences in relative expression levels among healthy tissues versus cancer cell lines (Fig. 5b), placenta versus other healthy tissues (Fig. 5c), and CML versus APL cell lines (Fig. 5d). These relative expression levels are consistent with previous literature reports of high expression of several miRNAs in placenta tissue^37^, higher expression of *hsa*-miR-141 in APL patients compared to CML patients with similar levels of *hsa*-miR-16 in both^38^, and high expression of *hsa*-miR-375 in lung tissue^39^. The relative expression levels of *hsa*-miR-375 and *hsa*-miR-16 across all tissue samples illustrate a dynamic range spanning from low-femtomolar to low-picomolar concentrations, allowing the simultaneous quantification of miRNAs with divergent expression levels (Fig. 5e).

**Fig. 5.**
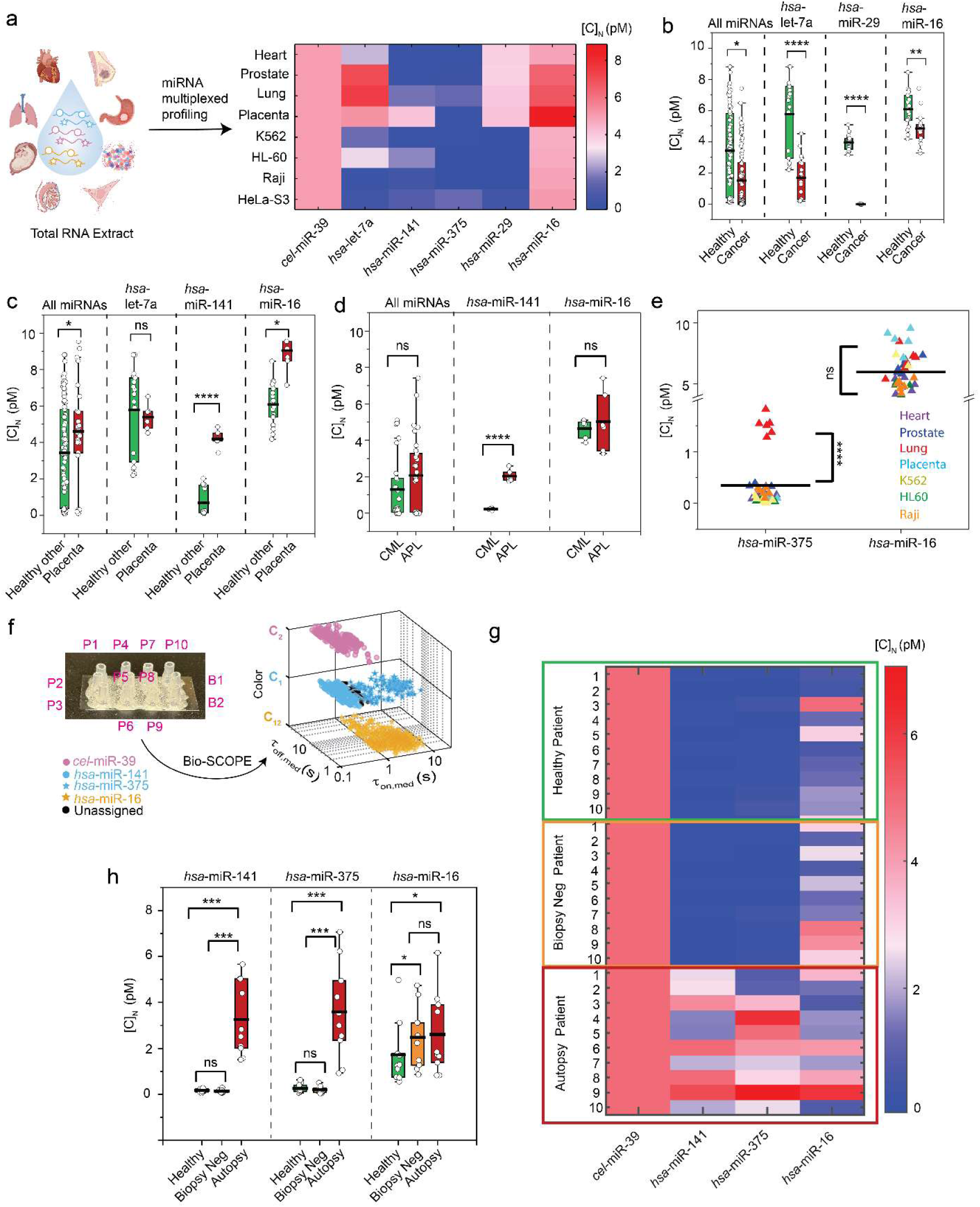
ǀ Multiplexed miRNA profiling of human tissues and prostate cancer patient samples using Bio-SCOPE. **a**, Heat map of the normalized concentrations [C]_N_ of six miRNAs (*hsa*-miR-141, *hsa*-miR-375, *cel*-miR-39, *hsa*-let-7a, *hsa*-miR-29, *hsa*-miR-16) as measured in multiplexed assays of total RNA extracts of eight different human tissue samples―four healthy tissues: heart, prostate, lung, placenta and four cancer tissues/cell lines: K562 (chronic myeloid leukemia, CML), HL-60 (acute promyelocytic leukemia, APL), Raji (Burkitt’s lymphoma), and HeLa-S3 (human cervical adenocarcinoma. *cel*-miR-39 was spiked into each sample at a known final concentration (i.e., 5 pM) for normalization. Relative expression levels of different miRNAs of interest in **b,** all healthy tissues versus cancer tissue samples **c,** all other healthy tissues versus placenta tissue samples, and **d,** CML versus APL samples. **e,** *hsa*-miR-375 and *hsa*-miR-16 relative expressions in all tissue samples. **f,** Multiplexed detection of two up-regulated (*hsa*-miR-141 and *hsa*-miR-375), one relatively unchanged (*hsa*-miR-16), and one spiked-in (*cel*-miR-39) miRNA in prostate cancer patient samples (total RNA extract from plasma) using Bio-SCOPE. Multiplexed miRNA profiling of one set of 10 patient samples can be done in a single coverslip with controls in other two chambers. Four miRNAs can be placed in 3D scatterplot of dwell time analysis (i.e., τ_on,med_ versus τ_off,med_) in different channel and kinetics with 95% confidence clusters (N_unassigned_ =16%). **g,** Heat map of [C]_N_ of miRNAs of interest in plasma from healthy (n=10), biopsy negative (n=10), and autopsy patient (n=10) samples. Similarly, *cel*-miR-39 is doped at known concentration into each patient sample to normalize the concentrations and minimize experimental errors. **h,** Relative expression levels of different miRNAs of interest in plasma from healthy (n=10), biopsy negative (n=10), and autopsy patient (n=10) samples. Data is presented as mean ± SD. Error bars indicate the SD of three independent measurements. Statistical significance was tested using two-tailed Student’s t-test (****P ≤ 10^-7^; ***P ≤ 10^-5^, **P ≤ 10^-3^, *P ≤ 10^-1^, not significant (ns) ≥ 0.1).

We next applied Bio-SCOPE to plasma samples from prostate cancer patients, detecting upregulated miRNAs (*hsa*-miR-141 and *hsa*-miR-375), one relatively unchanged miRNA (*hsa*-miR-16), and doped *cel*-miR-39 (for normalization) in total RNA extracts from plasma in a 4-plex assay (Fig. 5f)^4^. A heat map of the normalized concentrations [C]_N_ of the miRNAs in healthy controls (n=10), prostate biopsy-negative controls (n=10), and plasma samples collected at time of autopsy from patients with known prostate cancer (n=10) revealed distinct expression patterns (Fig. 5g). A two-tailed student’s T-test indicated significant differences in expression levels of *hsa*-miR-141 and *hsa*-miR-375, but not *hsa*-miR-16, among these groups (Fig. 5h), consistent with prior reports of elevated expression of *hsa*-miR-141 and *hsa*-miR-375, but not *hsa*-miR-16, in prostate cancer^4^. These findings demonstrate Bio-SCOPE’s capability for rapid, simultaneous multiplexed miRNA profiling with potential use in disease diagnostics and personalized medicine.

### Simultaneous detection of RNA, DNA and protein on a unified Bio-SCOPE platform

Recent evidence suggests that multiomic approaches provide superior diagnostics compared to single-modality biomarker detection, especially for a complex disease such as cancer^6,40–42^. To test Bio-SCOPE’s multiomics capability, we applied it to the simultaneous multiplexed detection of diverse types of biomarkers, including mutant DNAs, proteins, and miRNAs. First, we performed multiplexed detection of purified DNA mutants and proteins in buffer. Kinetic multiplexing of DNA fragments containing either the exon 19 del or the T790M DNA cancer mutation of the *EGFR* gene^43^ was achieved with clustering at 95% confidence. Additionally, multiplexed detection of the pro-inflammatory cytokines IL-8 and IL-6 was demonstrated following a protocol similar to that for miRNAs, but using an aptamer or Fab protein fragment previously developed as single-plex detection probes^30,31^ (Fig. 6a-c).

**Fig. 6.**
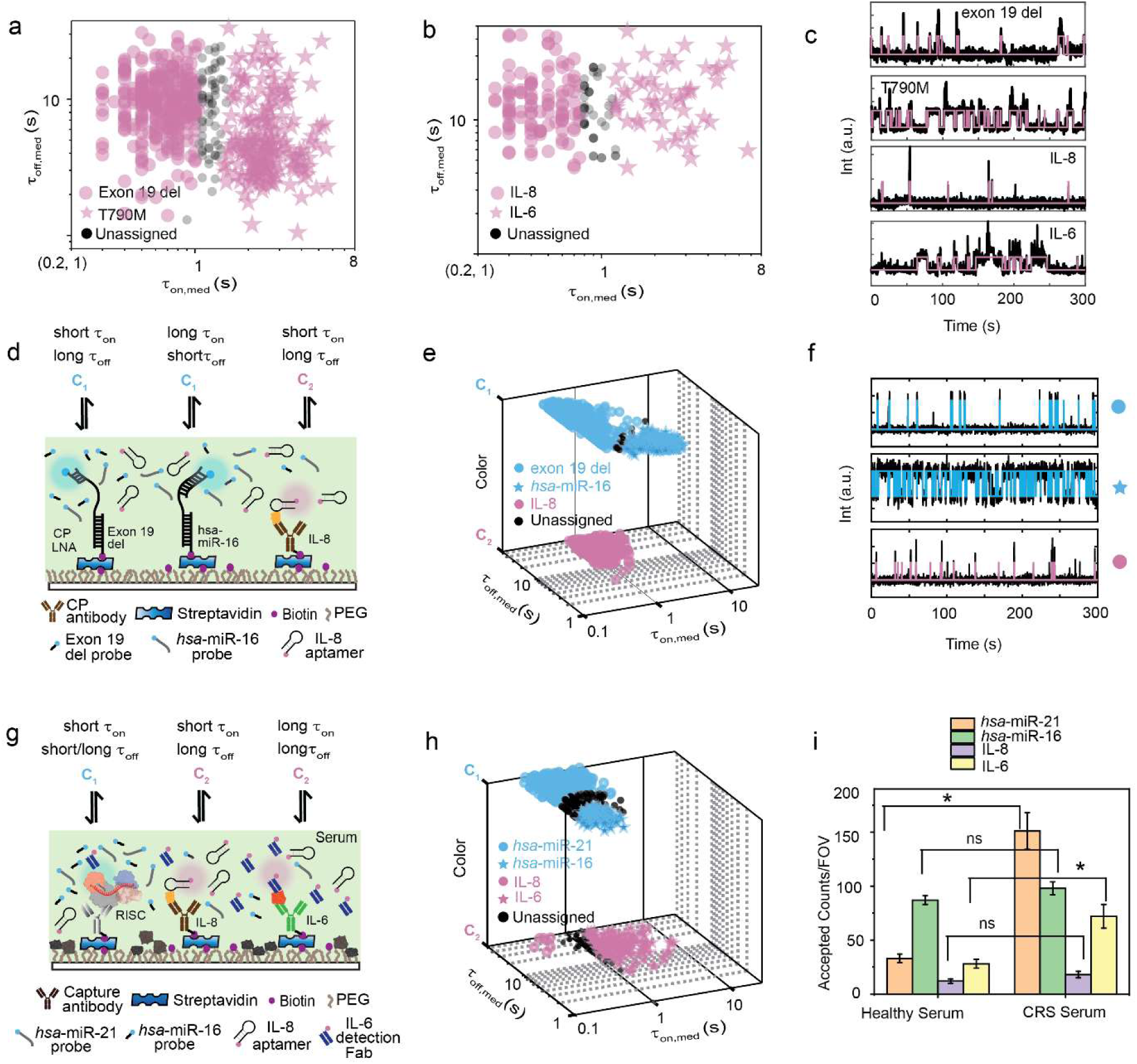
ǀ Multiplexed detection of miRNA, protein and DNA mutants. **a**, Kinetic multiplexing of EGFR exon 19 del and T790M mutants with 95% confidence clusters (N_unassigned_ =11%). **b,** Kinetic multiplexing of IL-8 and IL-6 with 95% confidence clusters (N_unassigned_ =13%). **c,** Representative single molecule trace behaviors of exon 19 del, T790M, IL-8, and IL-6. **d,** Schematic view of multiplexed detection of three types of biomarker molecules (miRNAs, mutant DNAs, and proteins) using Bio-SCOPE. **e,** 3D scatterplot of dwell time analysis (i.e., τ_on,med_ versus τ_off,med_ with color channels) for all accepted intensity-versus-time trajectories observed within single FOV of an equimolar mixture of exon 19 del (C_1_ channel, short τ_on,_ long τ_off_), *hsa*-miR-16 (C_1_ channel, long τ_on,_ short τ_off_), and IL-8 (C_2_ channel, short τ_on,_ long τ_off_; N_unassigned_ =2%) **f,** Representative single molecule trace behaviors of the above three targets. Unassigned molecules during the clustering are represented by black circles. Probes: exon 19 del _DNA_FP_Cy3_8-nt at 25 nM, *hsa*-miR-16_NS_DNA_FP_Cy3_C10-nt at 75 nM and IL-8_aptamer_8A-30_Cy5 at 50 nM. **g,** Schematic view of multiplexed detection of miRNAs and proteins in blood serum using Bio-SCOPE. **h,** 3D scatterplot of dwell time analysis (i.e., τ_on,med_ versus τ_off,med_ with color channels) for all intensity-versus-time trajectories observed within three FOV of multiplexed detection of miRNAs (N_unassigned_ =16%) and proteins (N_unassigned_ =18%) in CRS serum (with 95% confidence clusters). **i,** Multiplexed detection of different expressions of *hsa*-miR-21, *hsa*-miR-16, IL-8, and IL-6 in healthy control serum and in a serum sample from a cancer patient experiencing cytokine release syndrome (CRS) as a result of chimeric antigen receptor T cell (CAR-T cell) therapy. Black error bars represent the standard errors of the mean from two independent replicates. Statistical significance was tested using two-tailed Student’s t-test (*P ≤ 10^-1^, not significant (ns) ≥ 0.1). Serum concentration: 10%; Probes: *hsa*-miR-21_S_RNA_FP_Cy3_8-nt at 25 nM, *hsa*-miR-16_S_ RNA_FP_Cy3_7-nt at 75 nM, IL-8_aptamer_8A-30_Cy5 at 25 nM, IL-6_Fab_Cy5 at 75 nM.

Second, we developed a multi-omic assay simultaneously detecting three diverse biomarkers important in cancer diagnostics— exon 19 del DNA, *hsa*-miR-16 RNA, and IL-8 protein—which we tested by analyzing a synthetic, equimolar mixture of the three targets (Fig. 6d). All three biomarkers could be clustered by their distinct combinations of detection kinetics and color (Fig. 6e), as seen in representative time traces (Fig. 6f) and supported by cumulative dwell time analyses (Extended Data Fig. 6), demonstrating the power of chromato-kinetic encoding of the digital single molecule fingerprinting signal.

Third, we applied our multiplexed multiomic Bio-SCOPE assay directly to human blood serum samples. Notably, the direct single-molecule detection of biomarkers from unprocessed serum minimizes any variation or bias that may arise from multi-step pre-enrichment, extraction, purification, and/or amplification processes^7^. To achieve processing compatibility of blood-borne miRNA-loaded RISC detection^44^ with that of cytokines, we deployed single-molecule RISC pull-down using anti-Ago2 antibody^45,46^, followed by successful detection of specific miRNAs in serum using RNA probes that bind to the seed sequence of the miRNA target (Extended Data Fig. 7a). Based on this strategy, we achieved multiplexed detection of miRNAs *hsa*-miR-21 and *hsa*-miR-16 and cytokines IL-8 and IL-6 in human serum using a mixture of capture antibodies and detection probes of different types comprising oligonucleotides, an aptamer, and a Fab fragment (Fig. 6g, Extended Data Fig. 7b-h). *hsa*-miR-21 and *hsa*-miR-16 formed two distinct clusters in the C_1_ color channel, differing primarily based on τ_off_ (<τ_off_>_21_= 31.19 ± 0.08 s for miR-21, <τ_off_>_16_= 8.68 ± 0.02 s for miR-16), while IL-8 and IL-6 were identified as two distinct clusters in channel C_2_ based on τ_on_ (<τ_on_>_IL-8_= 0.27 ± 0.01 s, <τ_on_>_IL-6_= 2.27 ± 0.02 s) in the same multiplexing experiment.

Finally, we applied this Bio-SCOPE assay to the profiling of the relative expression levels of these four biomarkers in healthy serum versus serum from patients with diffuse large B-cell lymphoma (DLBCL) that received CAR-T cell therapy and experienced the common side effect of cytokine release syndrome (CRS)^47^. The assay revealed statistically significant upregulation of *hsa*-miR-21 and IL-6 in the patients with CRS, while the expression levels of *hsa*-miR-16 and IL-8 showed no such difference (Fig. 6i), consistent with the literature^30,48^. These results suggest that Bio-SCOPE can simultaneously profile both nucleic acid and protein biomarkers captured from crude biofluids without prior purification or workup.

## Discussion

We here demonstrate that Bio-SCOPE combines the unique analytical strengths of single-molecule kinetic fingerprinting and digital counting with multiplexing, opening up new opportunities for sensitive multiomic assays for molecular diagnostics. By exploiting unique kinetic fingerprints that distinguish specific targets from nonspecific interactions and contaminants, Bio-SCOPE not only achieves unrivaled specificity in single-molecule detection but expands beyond the typical limit of ∼3 colors for single-molecule detection through the added dimension of kinetics to permit simultaneous measurement of up to nine biomolecules. This digital encoding in the digital fingerprint is critical for applications such as molecular diagnostics, which often require the rapid, simultaneous measurement of multiple biomarkers. This multiplexing also facilitates normalization using internal controls, which we used to perform accurate miRNA expression profiling across different tissue types and disease states. Although multiplexing reduces the sensitivity somewhat compared to single-plex assays by reducing the surface density of capture probes for each analyte, the multiplexed assays retain low-femtomolar sensitivity and single-base mismatch selectivity, enabling accurate detection of even low-abundance targets.

Bio-SCOPE achieves its single-molecule sensitivity without any amplification, which confers two main benefits. First, it achieves similar performance for both nucleic acid and protein analytes, which promises combined high-sensitivity nucleic acid and protein (i.e., multiomic) panels on a unified platform. This is significant because integrating quantitative results for multiple divergent biomarkers can provide important information about physiology and pathology^6,40–42^. Bio-SCOPE’s ability to detect a range of molecular analytes may thus be useful in the identification of disease subpopulations and, ultimately, optimization of treatment strategies. Second, the direct amplification-free probing strategy renders Bio-SCOPE insensitive to PCR inhibitors, permitting its application to not only cell and tissue extracts but also crude human biofluids and potentially *in situ* measurements. We have herein demonstrated some of this potential of Bio-SCOPE by applying it to the quantitative, multiplexed profiling of miRNA, DNA mutations, and/or protein analytes from tissue and patient samples on a single analytical platform. Underscoring its potential utility in disease monitoring and personalized medicine, we used Bio-SCOPE to detect upregulation of specific miRNAs in prostate cancer samples, as well as elevated levels of cancer-related miRNA *hsa*-miR-21^32,33^ and pro-inflammatory cytokine IL-6 in CAR-T cell therapy patients with CRS^47^, providing insights into disease mechanisms and therapeutic responses.

Looking forward, Bio-SCOPE may be further enhanced by incorporating advanced machine learning algorithms for automatic trace classification, considering parameters such as signal intensity and signal-to-noise ratio^49^ to distinguish additional targets within a single-color channel, and thus further expanding multiplexing capabilities. Additionally, expanding the number of color channels, such as creating a channel where Cy3 and Cy5 are attached to the same probe, or using probes with variable ratios of emission in the two detection channels, will further broaden the method’s scope by digital encoding. Finally, Bio-SCOPE’s ability to provide comprehensive molecular profiles from minimal sample inputs—e.g., less than 1 µL with suitable microfluidics— could revolutionize clinical diagnostics by offering a holistic view of the molecular landscape, crucial for developing targeted therapies and personalized treatment plans. Overall, its unique combination of high-performance single-molecule detection, broad analyte scope, and multiplexing capabilities render Bio-SCOPE promising as a robust and versatile approach for research, quantitative disease diagnostics, and personalized medicine.

## Supporting information

Chromato-kinetic fingerprinting enables multiomic digital counting of single disease biomarker molecules

## Methods

### Oligonucleotides, proteins, and tissue RNA samples

All DNA and RNA oligonucleotides were purchased from Integrated DNA Technologies (IDT, www.idtdna.com) with high-performance liquid chromatography (HPLC) purification. miRCURY LNA miRNA power inhibitor (Catalog # 339131YI04102261-DDA) was obtained from Qiagen (www.qiagen.com). Sequences for all oligonucleotides are provided in Supporting Information (Supplementary Table 1). These oligonucleotides were dissolved in TE buffer (10 mM Tris, 0.1 mM EDTA, pH 8.0) to a final concentration of 100 µM, aliquoted and stored at –20°C for DNA or –80°C for RNA. Recombinant human IL-8 and IL-6, provided in lyophilized form by Bio-Rad Laboratories, Inc., were suspended in 2× PBS, pH 7.4 (Gibco) with 10 mg/mL BSA (Thermo Fisher Scientific Blocker BSA (10x) in PBS; catalog No. 37525) as a carrier, then aliquoted and stored at −80°C. Monoclonal IL-8 capture antibody (catalogue # EPR19358-108) and anti-Ago2 capture antibody (Catalog # ab57113) were purchased from Abcam. Monoclonal IL-6 capture antibody and detection Fab were used as previously reported^37^. All capture antibodies were bovine serum albumin (BSA) and azide-free to facilitate biotinylation using biotin NHS ester. Total RNA extracts from various human tissue samples were obtained from Thermo Fisher Scientific and stored at − 80°C after aliquoting. Human serum (catalogue # H4522) was purchased from Sigma Aldrich. All chemicals used for buffer preparation were obtained from Sigma-Aldrich or Thermo Fisher Scientific. For all experiments, DNA lo-bind tubes (Eppendorf) were used.

### Biotinylation of capture antibody

Monoclonal antibodies used for capture were biotinylated by attaching them to an amine-reactive NHS ester, specifically using biotin N-hydroxysuccinimidyl ester (Sigma Aldrich, Catalog #H1759-100) as described in previous studies^30,31^. For the reaction, a molar ratio of 5:1 for biotin to antibody was maintained at room temperature over 1 hour in 1× PBS with pH 7.4. Purification of the resulting biotin-IgG conjugates was achieved through the use of Zeba Spin desalting columns (Thermo Fisher, Catalog #89882, 7K MWCO) in accordance with the manufacturer’s instructions, followed by overnight dialysis at 4 °C against 1× PBS, pH 7.4 (Slide-A-Lyzer Dialysis Cassette, Thermo Fisher, 3.5K MWCO). The extent of biotinylation, estimated through an electrophoretic mobility shift assay with or without excess streptavidin, ranged between 70% and 80%. These capture antibodies were then aliquoted and stored at –80 °C.

### Fluorophore labeling of detection Fab fragments

Fab fragments for protein detection were labeled with a fluorescent dye via amine-NHS ester coupling using Cy5 mono-reactive dye packs (GE Healthcare, Catalog #PA25001). The labeling process used a molar ratio of 10:1 for dye to protein, performed in the dark at room temperature for 1 hour. Following the reaction, antibody-dye conjugates were purified with Zeba Spin desalting columns (Thermo Fisher, Catalog #89882, 7K MWCO) according to the manufacturer’s guidelines, and dialyzed overnight at 4°C against 1× PBS, pH 7.4 (Slide-A-Lyzer Dialysis Cassettes, Thermo Fisher, 3.5K MWCO). The dye to Fab fragment ratio following purification was verified to be 3:1 using UV-Vis spectrophotometry (Nanodrop) by measuring absorbance at 280 nm and 650 nm.

### Preparation of slide surfaces for single-molecule microscopy

Glass coverslips (No. 1.5, 24 × 50 mm, VWR Catalog #48393-241) were coated with a 1:100 mixture of biotin-PEG-SVA and mPEG-SVA (Laysan Bio, Inc., Catalog #MPEG-SVA-5000-1g and #BIO-PEG-SVA-5K-100MG) in accordance with previously established methods^36,37^. The modified coverslips were kept in a nitrogen-purged cabinet, wrapped in aluminum foil, for up to 4 weeks. Before conducting an experiment, 2-6 sample cells were prepared on each coverslip by cutting approximately 2 cm from the wider end of micropipette tips (Thermo Fisher, #02-682-261), discarding the narrower end, and placing the wide end down on the PEG-coated glass coverslip. The edges were then secured with epoxy adhesives (Ellsworth Adhesives, #4001).

### Single-molecule fluorescence microscopy

All single-molecule fluorescence experiments were performed using the Oxford Nanoimager (ONI), a compact benchtop microscope designed for objective-type total internal reflection fluorescence (TIRF) microscopy (Refer to https://oni.bio/nanoimager/ for camera, illumination, and objective specifications). The ONI featured a 100× 1.4NA oil-immersion objective and a Z-lock control module for maintaining autofocus. Since the ONI’s built-in temperature control system could not keep imaging temperatures below 25 °C, we prevented overheating by securing two cooling plates to the ONI’s outer box with metal clamps, allowing cold water from a water bath to flow through them. Prior to imaging, we pre-equilibrated the ONI to the desired temperature and adjusted the water bath temperature in real-time after laser activation to maintain a stable imaging temperature. All imaging was conducted at 25 °C. For TIRF, we utilized an illumination angle of approximately 54°. Cy3 and Cy5 fluorescence emissions were recorded with an optimal signal-to-noise ratio (S/N) by exciting the samples at 532 nm (at 36% laser power) and 640 nm (at 30% laser power), respectively (approximately 30 mW). The signal integration time per frame was set to 100 ms unless stated otherwise, with movies of 5 min duration collected per FOV. For experiments requiring chromatic multiplexing, movies were recorded on both channels simultaneously with both lasers active (Supplementary Fig. 2).

### Imaging solution

Unless stated otherwise, all Bio-SCOPE assays were conducted in an imaging solution that included 2× PBS, pH 7.4 (Gibco); an oxygen scavenger system^50^comprising 5 mM 3,4-dihydroxybenzoic acid (Fisher, #AC114891000), 0.05 mg/mL protocatechuate 3,4-dioxygenase (Sigma Aldrich, #P8279-25UN), and 1 mM Trolox (Fisher, #218940050); 1 µM LNA capture probe (CP) blocker; and 25-75 nM fluorophore-labeled oligonucleotide. In multiplexing experiments, multiple fluorophore-labeled oligonucleotides were introduced to the solution together, generally maintaining a combined concentration of up to 100 nM in each channel (Supplementary Fig. 4).

### Kinetic, chromatic, and chromato-kinetic fingerprinting assays

Sample cells were initially washed with 100 µL T50 buffer (10 mM Tris-HCl, pH 8.0 at 25°C, 50 mM NaCl) and then incubated with 40 µL of a 1 mg/mL streptavidin solution in T50 buffer for 10 min. After removing excess streptavidin, the cells were washed three times with 100 µL of 2× PBS. The coverslip was then coated with the biotinylated capture probe (CP; LNA or antibody) by adding 40 µL of a 100 nM biotinylated CP solution in 2× PBS and incubating for 10 to 30 min. Excess CP was removed, and the sample wells were washed three times with 100 µL of 2× PBS. For multiplexing experiments, multiple CPs were incubated simultaneously, keeping the total concentration at 100 nM. Typically, the number of CPs corresponded to the number of targets unless specified otherwise. The target or target mixture (100 µL) in 2× PBS was then incubated for 1 hour at room temperature. For tissue sample experiments, RNA extracts were diluted 100-fold in 2× PBS (Supplementary Fig. 6), and for patient samples, the dilution was 2-fold, with the final target volume being 20 µL in both cases. After capturing the target, the sample cells were washed three times with 2× PBS, and then 100 µL of imaging solution was added. Samples were immediately imaged using TIRF microscopy.

### Analysis of Bio-SCOPE data, *k*-means clustering, and 3D scatter plot

Bio-SCOPE data were analyzed using custom MATLAB scripts to identify fluorophore-labeled oligonucleotide probe binding sites and examine the kinetics of repeated binding, as described previously with a diffraction-limited analysis method^28^ (Supplementary Fig. 2). Briefly, regions of repeated probe binding and dissociation, designated as regions of interest (ROIs) within the FOV, were identified by calculating the average absolute frame-to-frame intensity change at each pixel to create an intensity fluctuation map. ROIs were then defined as 3×3-pixel regions centered on local maxima within this fluctuation map. The integrated, background-subtracted intensity within each ROI was subsequently calculated for each movie frame to generate an intensity-versus-time trace. These candidate traces were subjected to hidden Markov modeling (HMM) using a customized version of vbFRET^28^. The idealized trace produced by HMM was used to determine several parameters for fingerprinting analysis: N_b+d_, the number of binding and dissociation events; τ_on,med_ and τ_off,med_ the median dwell times in the probe-bound and probe-unbound states, respectively; τ_off,max_, the maximum dwell time in the probe-unbound state; and S/N, the signal-to-noise ratio, defined as the standard deviation of the fluorescence intensity divided by the mean intensity difference between bound and unbound states (Supplementary Table 2). Threshold values for each of these parameters to count a trace as a positive detection event were optimized using a custom optimizer software for each detection probe. The C_12_ channel was formulated as described earlier in the main text and in Supplementary Fig. 2.

To determine target identity in a mixture of targets, *k*-means clustering was used to classify dwell times (i.e., τ_on,med_ and τ_off,med_) from each dataset containing multiple targets. The mean and standard deviation for each cluster were calculated using a 3D Gaussian fit without covariance to the cluster, after discarding all outliers (>3σ from the initial cluster mean). Each cluster was pretrained using data from separate individual experiments conducted under the same conditions (Fig. 2b). This clustering method enabled the identification of at least three targets within a single-color channel (Fig. 2b). Additionally, combining three channels (i.e., C_1_, C_2_, and C_12_) allowed for the concurrent identification of multiple targets, which could then be represented by a 3D scatter plot using color as the third dimension (Supplementary Fig. 4).

### Fitting of the cumulative frequency of dwell times

Average dwell times for each experiment were analyzed using a custom MATLAB script. The script initially calculated the cumulative frequency of all dwell times for a given state using bins corresponding to the camera exposure time (100 ms). This cumulative frequency was then fitted to an exponential function (Equation 1):

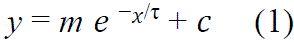

where *m*, *n*, *c*, and τ are fit parameters for S1. For the single exponential fit, the coefficient m is set to –1. The coefficient τ represents the average dwell time for a given event in the single exponential fit. The coefficient c is a constant indicating the y-intercept of the equation. Each dataset’s cumulative frequency was fitted to the single exponential function. This fit was accepted if the sum squared error was less than 0.05 and R^2^ was greater than 0.99, indicating a good fit and suggesting that the coefficient τ accurately represented the average dwell time.

### Limit of detection (LOD) estimation for individual and multiplexing experiments

Bio-SCOPE of FP binding was imaged by TIRF microscopy using an acquisition time of 5 min / FOV at 25°C with varying concentrations of synthetic miRNAs (Fig. 1b). Standard curves were then analyzed by fitting a linear regression model. LOD was estimated by dividing three times the standard deviation of blanks by the slope of the standard curve, as follows:

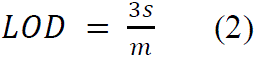

where *s* is the standard deviation of blank and m is the slope of the standard curve. LOD estimation of hsa-miR-16 in multiplexing scenarios was performed similarly, but in the presence of five other miRNAs at constant concentrations (Fig. 3b).

### Blood sample collection

Healthy control blood was collected under University of Michigan IRB protocol HUM00092161. Plasma was collected in an EDTA vacutainer, and spun at 1,600 × g for 10 min. The plasma was then transferred to a new tube and spun at 16,000 × g for 10 min. The plasma was subsequently transferred to a new tube and frozen at –80 °C. Serum samples were collected without an anticoagulant, allowed to clot, and spun at 1,200 × g for 15 min. The serum was transferred to a new tube and frozen at –80 °C. CRS patient serum samples were collected and handled as previously reported^30^.

Serum was collected from consented patients according to the Prostate Biopsy Clinical Database/Tissue Bank Protocol at the University of Michigan. It was approved by the Institutional Review Board and Ethics Committee and supported by the National Cancer Institute (NCI) Early Detection Research Network (EDRN). Blood was centrifuged for 20 minutes at 1300g-force, 4°C. Serum was aliquoted and stored at –70°C or colder. Clinical information obtained for each patient included age, race, serum PSA, family history, pathology, results, medical history/ comorbidities, and follow up data.

EDTA plasma was collected from the rapid autopsy program approved by the Institutional Review Board of the University of Michigan and supported by the NCI Specialized Program of Research Excellence in Prostate Cancer (SPORE). Clinical information obtained for each patient included age at diagnosis, age at death, months to death after diagnosis, initial Gleason score, time on hormonal therapy only, and type of primary and systemic therapy. EDTA blood was centrifuged for 20 minutes at 1300g-force, 4°C. Plasma was aliquoted and stored at –70°C or colder.

### RNA extraction from plasma and serum

RNA was extracted from plasma and serum samples using the miRNeasy Mini Kit (Qiagen) following the manufacturer’s protocol with minor adjustments. Specifically, 200 µL of the sample was combined with 1 mL of Qiazol, vortexed, and then incubated at room temperature for 5 min. Afterward, 200 µL of chloroform was added, vortexed again, and left at room temperature for 2 min. The samples were then centrifuged for 15 min at 12,000 × g at 4 °C. The aqueous phase was carefully transferred to a new tube, mixed with 1.5× volumes of 100% ethanol, and processed using the provided columns and buffers. The isolated RNA was spiked with *cel*-miR-39 to a final concentration of 5 pM.

### Bio-SCOPE for multiplexed multiomic detection of endogenous targets in serum

To detect endogenous miRNAs in serum, the RISC was captured using biotinylated anti-Ago2 capture antibodies (10 nM concentration, 1-h incubation) following a previously reported protocol^45–46^. Endogenous *hsa*-miR-21 and *hsa*-miR-16 were then simultaneously detected in 10% serum using fluorophore-labeled RNA oligonucleotides designed to bind the seed sequence of the miRNA. Additionally, endogenous IL-8 and IL-6 were captured using their respective CP antibodies and detected via aptamers (for IL-8) or detection Fab fragment (for IL-6). These assays were performed simultaneously in serum using a mixture of CP antibodies and detection FPs through the Bio-SCOPE assay (Fig. 6g-i).

## Data availability

The data supporting the findings of this study are available within this Article and its Supplementary Information. Source data are provided with this paper. Any additional data are available from the corresponding author upon request.

## Code availability

The code used in this study has been described in reference^28–31^ and is available through a DeepBlue repository. Any additional information concerning the code is available from the corresponding author upon request.

## Acknowledgements

We thank Professor Arul Chinnaiyan (Director, Michigan Center for Translational Pathology, University of Michigan) for providing clinical samples from prostate cancer patients and Professor Sung Won Choi for assistance in obtaining blood samples from patients receiving CAR-T cell therapy. We thank Alexander Johnson-Buck for his invaluable proofreading and feedback on the manuscript. This work was supported by NIH grants R21 CA225493 (N.G.W. and M.T.) and MIRA R35 GM131922 (N.G.W.). This work was additionally supported by the NCI EDRN (grant number U2C CA271854 to A.M.C.) and the NCI SPORE program (grant number P50 CA186786 to A.M.C.).

## Author contributions

P.B., S.R, S.M. and N.G.W. conceived the idea. P.B. and S.R designed experiments. P.B. performed most experiments. P.B., S.R. and L.D. analyzed, discussed the data and created the figures. E.S. and J.S. collected and prepared human plasma samples and performed RNA extraction. T.C. developed the miRISC based assay. P.B. wrote the initial draft of the manuscript. M.T. and N.G.W. were responsible for the conception and supervision of the project. All authors reviewed and edited the manuscript.

## Additional information

**Supplementary information** Supplementary information is available in the online version of the paper.

**Correspondence and requests for materials** should be addressed to Nils G. Walter.

**Extended Data Fig. 1.**
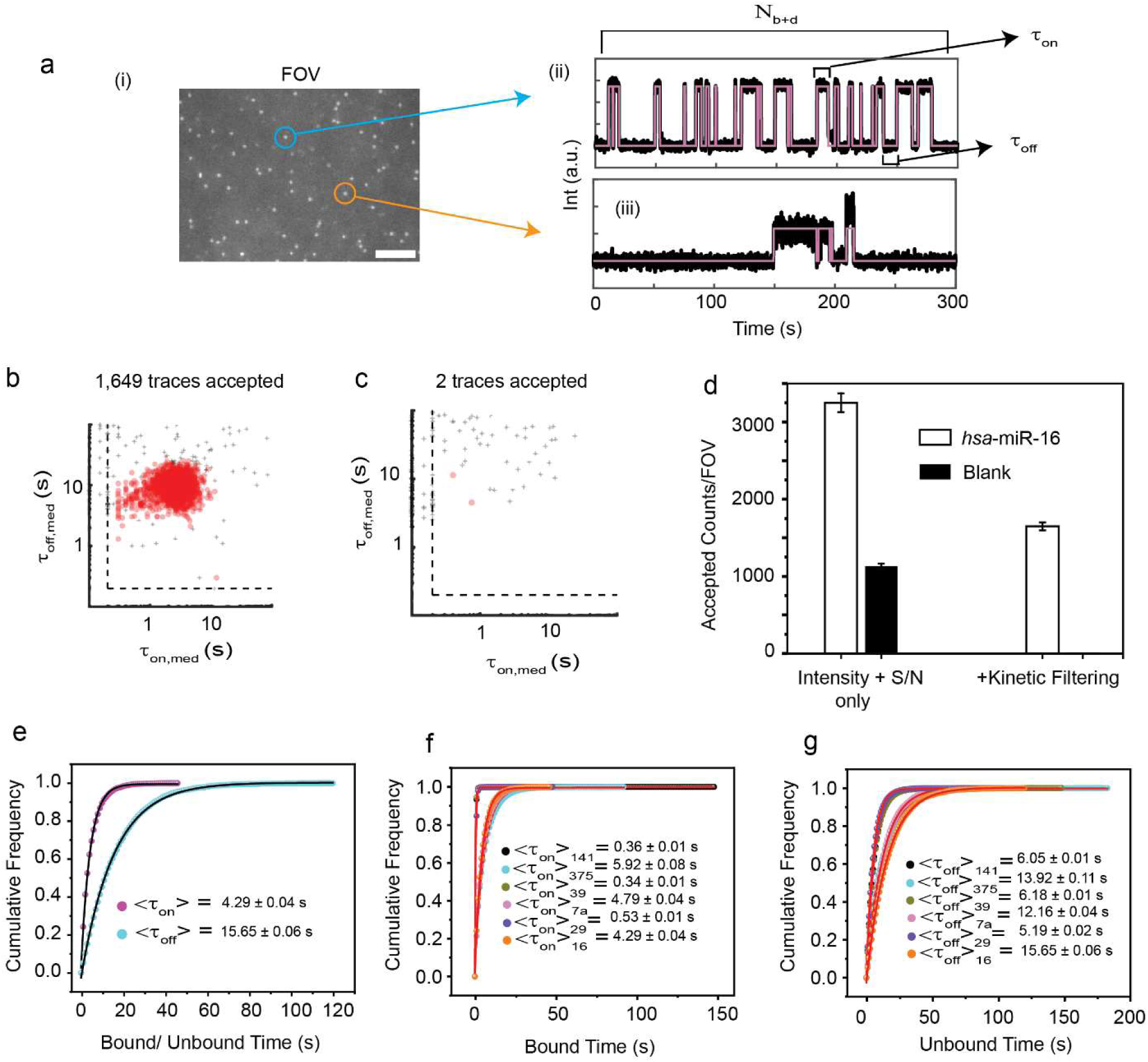
ǀ High-confidence detection of specific targets by kinetic fingerprinting. **a**, (i) Single movie frame of a representative portion of a Total Internal Reflection Fluorescence (TIRF) microscope field of view (FOV; scale bar: 5 µm), showing bright puncta at the locations where single fluorescent probes are bound at or near the imaging surface. (ii, iii) Representative kinetic fingerprints indicative of repetitive fluorescent probe binding to the same target molecule (here i.e., *hsa*-miR-16) (top) and nonspecific binding, which is typically less repetitive (bottom). Raw intensity-versus-time traces (black lines) are idealized by hidden Markov modeling (magenta lines) to extract kinetic parameters for analysis. Bound time (τ_on_), unbound time (τ_off_) and number of binding and dissociation events (N_b+d_) of a single trajectory are indicated by arrows in (ii). (**b** and **c**) Scatterplots of dwell time analysis (i.e., τ_on,med_ versus τ_off,med_) for all intensity-versus-time trajectories observed within a single field of view in the presence (b) or absence (c) of 5 pM *hsa*-miR-16. Dashed lines indicate thresholds (minimum or maximum) for accepting a trajectory as evidence of a single *hsa*-miR-16 molecules. Points indicated by “+” represent trajectories that do not pass filtering for intensity, signal-to-noise ratio, and/or kinetics, and are not considered sufficient evidence to detect *hsa*-miR-16. Points indicated by red-filled circles represent trajectories that pass filtering and are considered positive detection events of single hsa-miR-16 molecules. Temperature: 25°C; Acquisition time: 5 min/FOV; Probe: *hsa*-miR-16_NS_DNA_FP_Cy5_C10-nt at 50 nM. **d,** Impact of kinetic filtering on the number of accepted *hsa*-miR-16 counts in the presence and absence of *hsa*-miR-16. Black error bars represent the standard errors of the mean from three independent replicates. **e,** Cumulative bound (magenta circles) and unbound (cyan circles) dwell time histograms of Cy5-labelled *hsa*-miR-16 (*hsa*-miR-16_NS_DNA_FP_Cy5_C10-nt at 50 nM). <τ_on_> and <τ_off_> values were determined from single-exponential fitting of cumulative dwell time distributions derived from 1,649 accepted single-molecule traces. Cumulative (**f**) bound and (**g**) unbound dwell time histograms of target specific NS DNA FPs with six individual optimized miRNAs (i.e., *hsa*-miR-141, *hsa*-miR-375, *cel*-miR-39, *hsa*-let-7a, *hsa*-miR-29, *hsa*-miR-16). The <τ_on_> and <τ_off_> values were determined from single-exponential fitting of cumulative dwell time distributions of accepted single-molecule traces. Target concentration: 5 pM; Temperature: 25°C; Acquisition time: 5 min/FOV, Probes: *hsa*-miR-141_NS_DNA_ FP_Cy3_8-nt at 50 nM, *hsa*-miR-375_NS_DNA_FP_Cy3_C8-nt at 50 nM, *cel*-miR-39_NS_DNA_ FP_Cy5_8-nt at 50 nM, *hsa*-let-7a_NS_DNA_FP_Cy5_11-nt at 50 nM, *hsa*-miR-29_NS_DNA_ FP_Cy3 & Cy5_8-nt at 50 nM, *hsa*-miR-16_NS_DNA_FP_Cy3 & Cy5_C10-nt at 50 nM.

**Extended Data Fig. 2.**
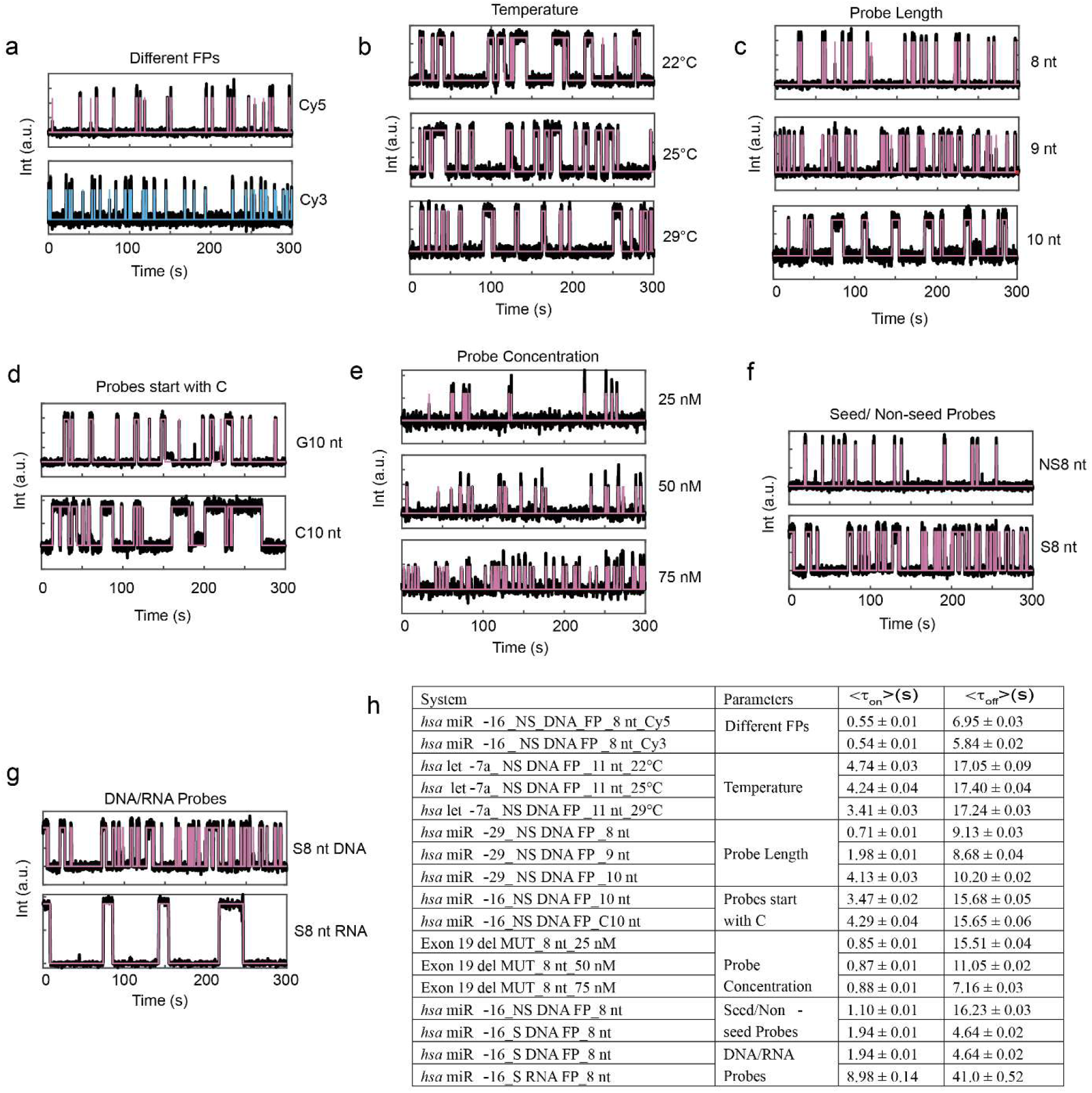
ǀ Generating diverse kinetic fingerprinting using different experimental parameters. Representative kinetic fingerprinting (i.e., single-molecule intensity-versus-time traces from TIRF microscopy measurements) of **a**, Different FPs: *hsa*-miR-16 with NS_DNA_FP_8-nt_Cy5 or _Cy3 at 50 nM, 25°C. **b,** Temperature: *hsa*-let-7a with NS_DNA_FP_11-nt_Cy5 at 50 nM and 22°C, 25°C or 29°C. **c,** Probe length: *hsa*-miR-29 with NS_ DNA_FP_Cy5_8-nt, _9-nt or _10-nt at 50 nM and 25°C. **d,** Probes starting with C: *hsa*-miR-16 with NS_DNA_FP_Cy5_G10-nt and C10-nt at 50 nM dye concentration and 25°C. **e,** Probe concentration: exon 19 del MUT with DNA_FP_Cy5_8-nt at 25 nM, 50 nM and 75 nM and 25°C. **f,** Seed/Non-seed probes: *hsa*-miR-16 with DNA_FP_Cy5_8-nt, seed and non-seed probes at 50 nM and 25°C. **g,** DNA/RNA probes: *hsa*-miR-16 with S_ FP_Cy5_8-nt, DNA and RNA probes at 50 nM and 25°C. **h,** Table of different <τ_on_> and <τ_off_> values obtained from single-exponential fitting of cumulative dwell time distributions in different scenarios. The target concentration was 5 pM throughout.

**Extended Data Fig. 3.**
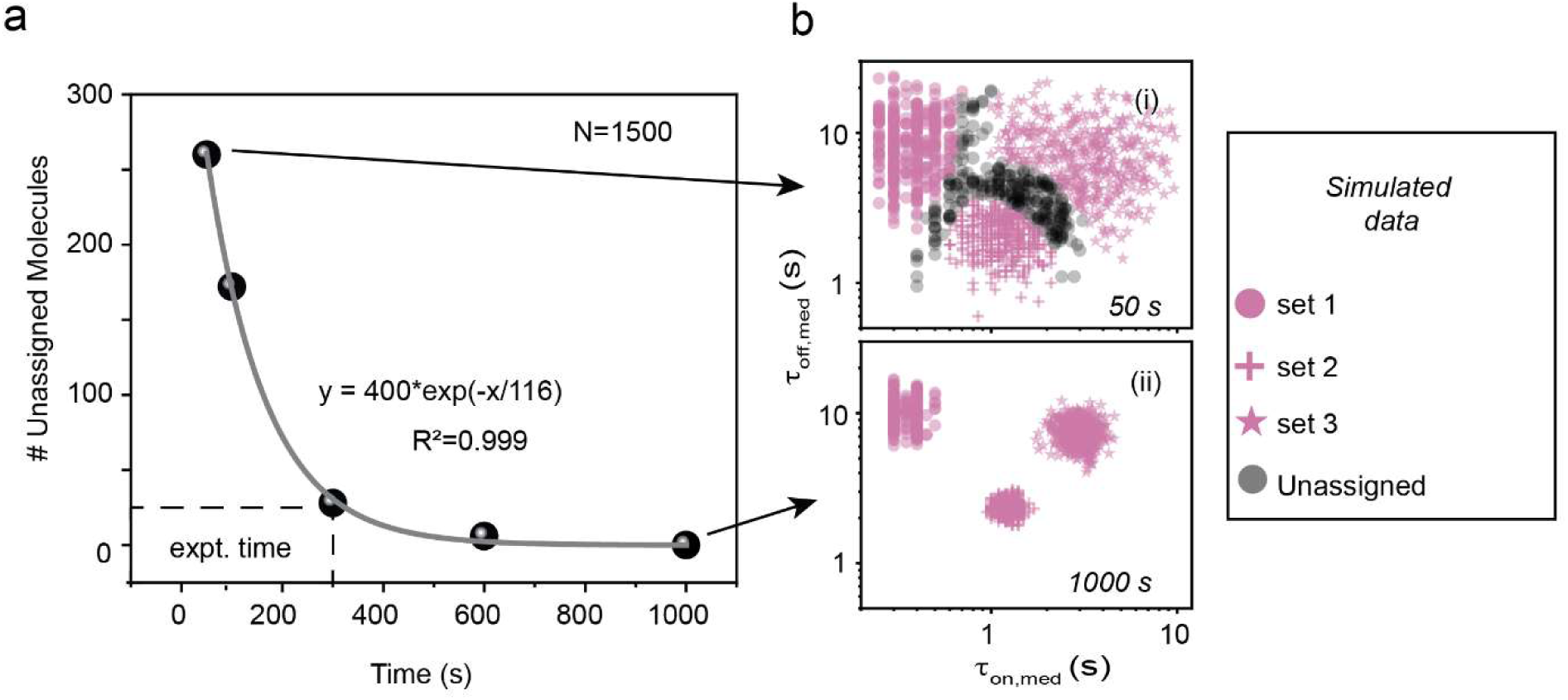
ǀ Clustering of kinetic multiplexing in simulated trace datasets. **a**, Number of unassigned molecules in 95% confidence clusters with increasing acquisition time for simulated trace datasets (total number of molecules, N=1500). The dotted line indicates the number of unassigned molecules at the experimental acquisition time (i.e., 300 s or 5 min). **b,** Kinetic multiplexing with 95% confidence clusters for three different targets in simulated trace datasets with acquisition times of (i) 50 s (N_unassigned_ = 17%) and (ii) 1000 s (N_unassigned_ = 0%).

**Extended Data Fig. 4.**
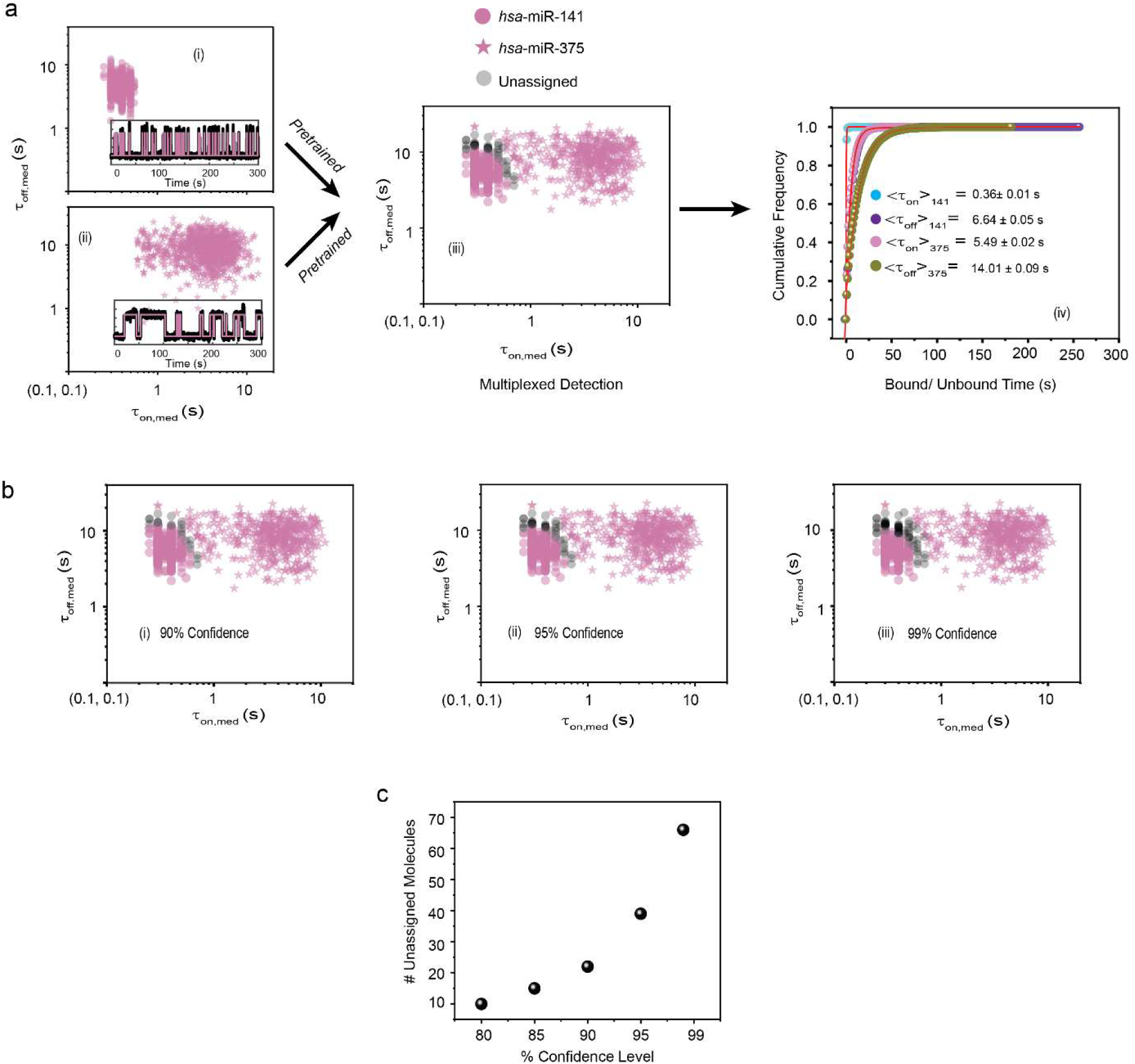
ǀ Clustering of kinetic multiplexing datasets (two-plex). **a**, Scatterplots of dwell time analysis (i.e., τ_on,med_ versus τ_off,med_) for all accepted intensity-versus-time trajectories observed within a single field of view of (i) *hsa*-miR-141, (ii) *hsa*-miR-375 and (iii) a 1:1 mixture of *hsa*-miR-141 and *hsa*-miR-375 (with 95% confidence clusters, N_unassigned_ =4%). Unassigned molecules during two-plex clustering are represented with black circles. Representative single molecule trace behaviors are shown in the insets. (iv) Cumulative bound and unbound dwell time histograms of *hsa*-miR-141 and *hsa*-miR-375. **b,** Kinetic multiplexing of the two-plex dataset with (i) 90% (N_unassigned_ =2%), (ii) 95% (N_unassigned_ =4%), and (iii) 99% (N_unassigned_ =6%) confidence clusters. **c,** Number of unassigned molecules as a function of confidence levels. Target concentration: 5 pM; Temperature: 25°C; Acquisition time: 5 min/FOV; Probes: *hsa*-miR-141_NS_DNA_FP_Cy3_8-nt at 50 nM, *hsa*-miR-375_NS_DNA_FP_Cy3_C8-nt at 50 nM.

**Extended Data Fig. 5.**
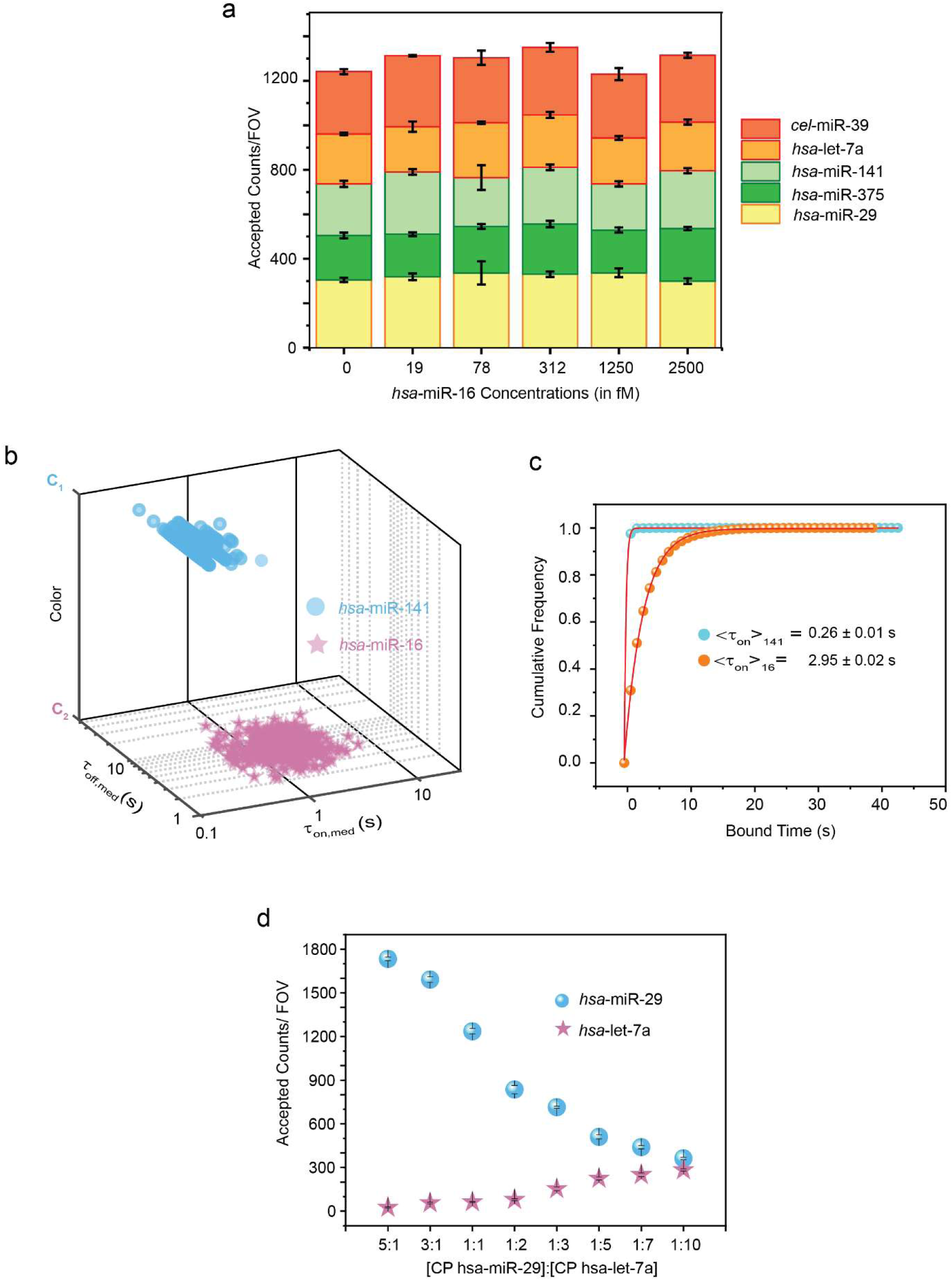
ǀ Sensitivity and multiplexing using Bio-SCOPE for different ratio detection. **a**, Change in accepted counts per field of view (FOV) of five other miRNAs (i.e., *cel*-miR-39, *hsa*-let-7a, *hsa*-miR-141, *hsa*-miR-375, *hsa*-miR-29) during *hsa*-miR-16 LOD estimation in multiplexing scenarios. Black error bars represent the standard errors of the mean from three independent replicates. **b,** 3D scatterplot of dwell time analysis (i.e., τ_on,med_ versus τ_off,med_ with color channels) for all accepted intensity-versus-time trajectories observed within single field of view of an equimolar mixture of *hsa*-miR-141 (Cy3 channel, fast kinetics) and *hsa*-miR-16 (Cy5 channel, slow kinetics). **c,** Cumulative bound time histograms of *hsa*-miR-141 and *hsa*-miR-16. Temperature: 25°C; Acquisition time: 5 min/FOV; Probes: *hsa*-miR-141_NS_DNA_FP_Cy3_8-nt at 50 nM, *hsa*-miR-16_NS_DNA_FP_Cy5_C10-nt at 50 nM. **d,** Detection of *hsa*-miR-29 (high abundance, Cy5 channel, fast kinetics) and *hsa*-let-7a (low abundance, Cy3 channel, slow kinetics) in a 1:1 ratio by varying capture probe concentrations. Black error bars represent the standard errors of the mean from three independent replicates. Total target concentration: 5 pM (1:1); Temperature: 25°C; Acquisition time: 5 min/FOV; Probes: *hsa*-miR-29_NS_DNA_FP_Cy5_8-nt at 50 nM, *hsa*-let-7a_NS_DNA_FP_Cy3_11-nt at 50 nM.

**Extended Data Fig. 6.**
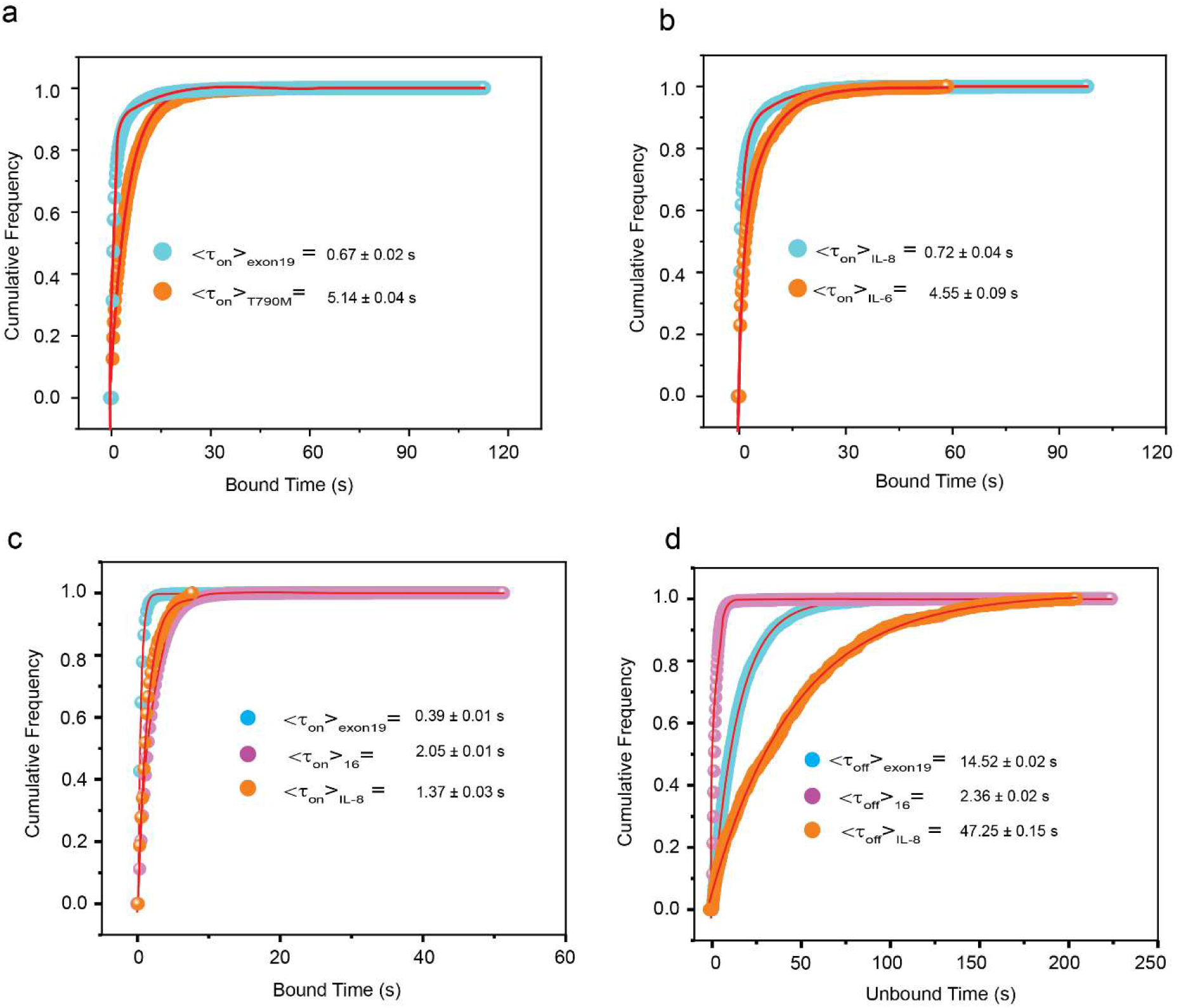
ǀ Cumulative dwell time analysis of targets in multiplexed detection of DNA MUTs, proteins and multiomic detection by different kinetic fingerprinting. **a**, Cumulative bound time histograms of (i) exon 19 del and (ii) T790M. <τ_on_> values were determined from single-exponential fitting of cumulative bound time distributions of accepted single-molecule traces. **b,** Cumulative bound time histograms of IL-8 and IL-6. <τ_on_> values were determined from single-exponential fitting of cumulative bound time distributions of accepted single-molecule traces. Total target concentrations: 5 pM (1:1); Temperature: 25°C; Acquisition time: 5 min/FOV; Probes: exon 19 del _DNA_FP_Cy5_8-nt at 25 nM, T790M_DNA_FP_Cy5_G9-nt at 75 nM, IL-8_aptamer_8A-30_Cy5 at 25 nM, IL-6_Fab_Cy5 at 75 nM. Cumulative **(c)** bound and **(d)** unbound dwell time histograms of exon 19 del, *hsa*-miR-16 and IL-8 for multiomic detection. <τ_on_> and <τ_off_> values were determined from single-exponential fitting of cumulative dwell time distributions of accepted single-molecule traces. Temperature: 25°C; Acquisition time: 5 min/FOV; Probes: exon 19 del_DNA_FP_Cy3_8-nt at 25 nM, *hsa*-miR-16_NS_DNA_FP_Cy3_C10-nt at 75 nM and IL-8_aptamer_8A-30_Cy5 at 50 nM.

**Extended Data Fig. 7.**
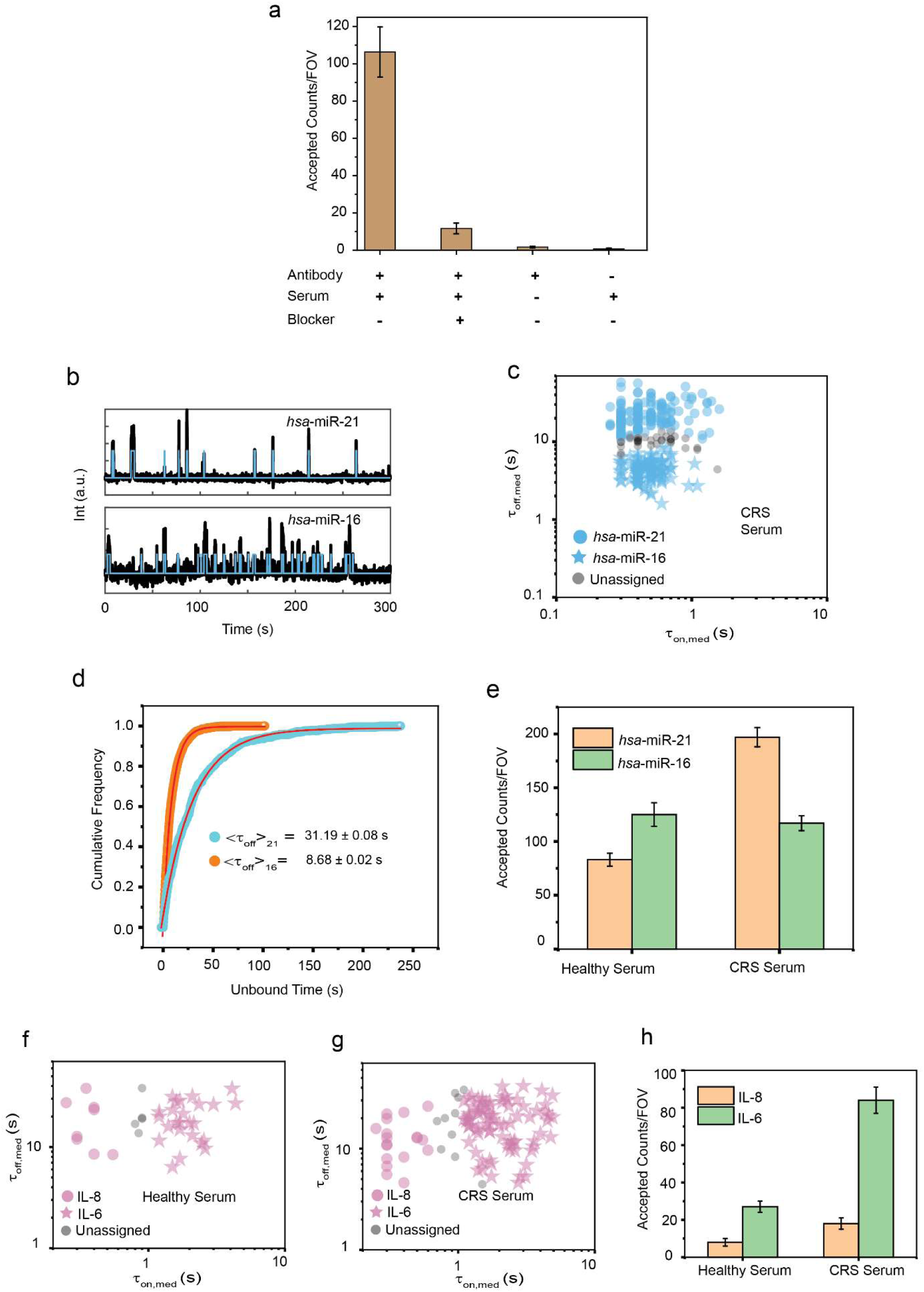
ǀ Multiplexed detection of miRNAs and proteins in serum. **a**, The number of accepted counts/FOV under different conditions. Kinetic filtering keeps the accepted counts near zero in the absence of either serum or capture antibody. Addition of *hsa*-miR-21 blocker lowers the accepted counts significantly by blocking access to *hsa*-miR-21 by fluorescent probes. Black error bars represent the standard errors of the mean from three independent replicates. **b,** Representative kinetic fingerprinting (i.e., single-molecule intensity-versus-time traces from TIRF microscopy measurements) of *hsa*-miR-21 and *hsa*-miR-16 detected in CRS serum. **c,** Scatterplots of dwell time analysis (i.e., τ_on,med_ versus τ_off,med_) for all intensity-versus-time trajectories observed within a single field of view of multiplexed detection of *hsa*-miR-21 and *hsa*-miR-16 in CRS serum (with 95% confidence clusters, N_unassigned_ =9%). **d,** Cumulative unbound time histograms of (i) *hsa*-miR-21 and (ii) *hsa*-miR-16. <τ_off_> values were determined from single-exponential fitting of cumulative bound time distributions of accepted single-molecule traces. **e,** Different expressions of *hsa*-miR-21 and *hsa*-miR-16 in healthy serum and CRS serum in multiplexing scenarios. Black error bars represent the standard errors of the mean from three independent replicates. Serum concentration: 10%; Temperature: 25°C; Acquisition time: 5 min/FOV; Probes: *hsa*-miR-21_S_RNA_FP_Cy3_8-nt at 25 nM, *hsa*-miR-16_S_RNA_FP_Cy3_7-nt at 75 nM.n**f,** Scatterplots of dwell time analysis (i.e., τ_on,med_ versus τ_off,med_) for all intensity-versus-time trajectories observed within a single field of view of multiplexed detection of IL-8 and IL-6 in (a) healthy serum (N_unassigned_ =17%) and (b) CRS serum (N_unassigned_ =11%) (with 95% confidence clusters). **g,** Different expressions of IL-8 and IL-6 in healthy serum and CRS serum in multiplexing scenarios. Black error bars represent the standard errors of the mean from three independent replicates. Serum concentration: 10%; Temperature: 25°C; Acquisition time: 5 min/FOV; Probes: IL-8_aptamer_8A-30_Cy5 at 25 nM, IL-6_Fab_Cy5 at 75 nM.

