## Supplementary material for "Chromato-kinetic fingerprinting enables multiomic digital counting of single disease biomarker molecules": Chromato-kinetic fingerprinting enables multiomic digital counting of single disease biomarker molecules

### Table of Contents

Table of Contents ……………………………………………………………………….…………2

Supplementary Figures…………………………………………………………….………....3

Supplementary Fig. 1 | Dwell times and clustering of individual two-plex datasets for kinetic multiplexing three-plex…………………………………………...…………………….…3

Supplementary Fig. 2 | Pipeline of Bio-SCOPE with three-color channels……………………………………………………………………………..….4

Supplementary Fig. 3 | Cumulative dwell time analysis of chromatic multiplexing three-plex………………………………………………………………………………….…..5

Supplementary Fig. 4 | Pipeline of Bio-SCOPE…………………………………………....6

Supplementary Fig. 5 | Bound times and clustering with different confidence levels of let-7 family three-plex datasets ………………………………………….………………......7

Supplementary Fig. 6 | Tissue Sample dilution and *cel*-miR-39 doping……………….……….... 9

Supplementary Tables…………………………………………………………….……...10

Supplementary Table 1| miRNA/DNA/Protein sample names and sequences……………………10

Supplementary Table 2 | Optimized general parameter sets for trace generation and analysis…....12

### Supplementary Figures

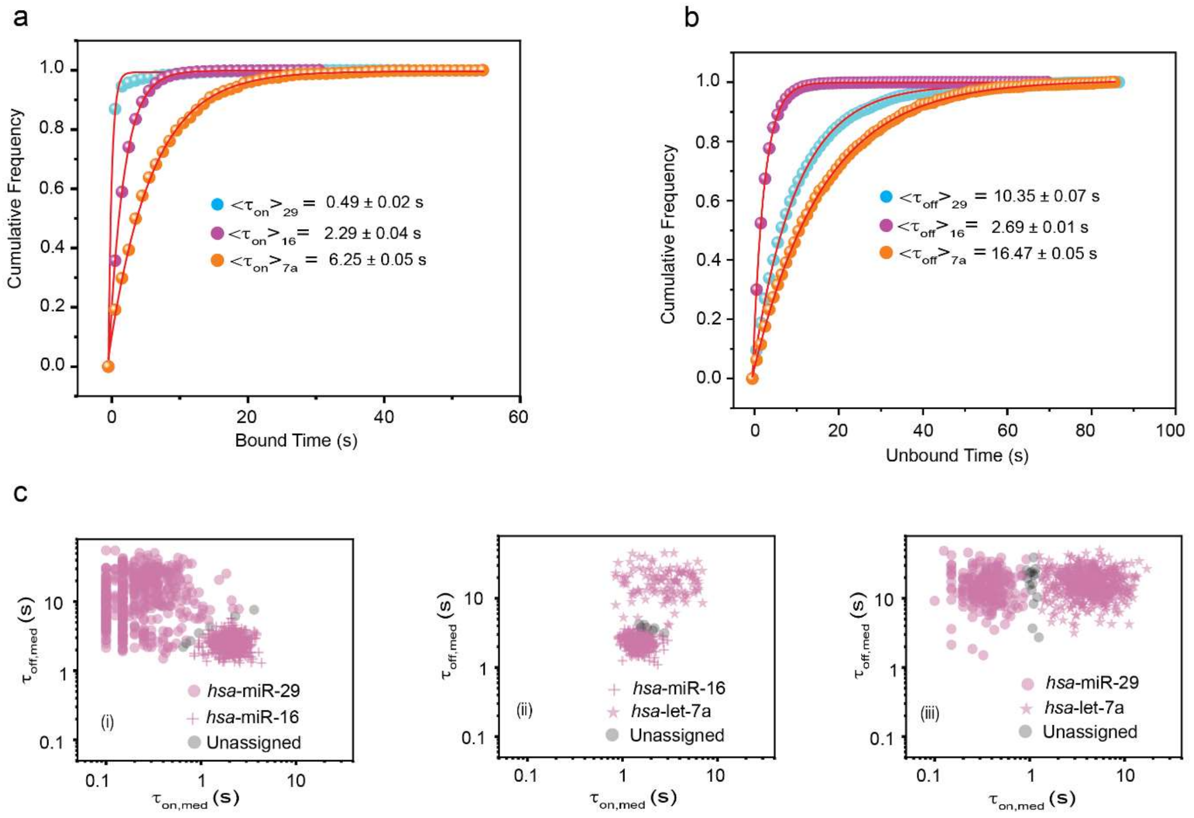


**Supplementary Fig. 1 | Dwell times and clustering of individual two-plex datasets for kinetics-based three-plex assay.** Cumulative (**a**) bound and (**b**) unbound dwell time histograms of *hsa*-miR-29, *hsa*-miR-16 and *hsa*-let-7a for kinetic multiplexing three-plex assay. $\langle\tau_{on}\rangle$ and $\langle\tau_{off}\rangle$ values were determined from single-exponential fitting of cumulative dwell time distributions of accepted single-molecule traces. **c,** Kinetic multiplexing of the individual two-plex datasets with 95% confidence clusters to achieve three-plex: (i) *hsa*-miR-29 and *hsa*-miR-16 (1:1) ($N_{29}$ = 51%, $N_{16}$ = 48%, $N_{unassigned}$ = 1%), (ii) *hsa*-miR-16 and *hsa*-let-7a (1:5) ($N_{16}$ = 64%, $N_{7a}$ = 35%, $N_{unassigned}$ = 1%) and (iii) *hsa*-miR-29 and *hsa*-let-7a (1:5) ($N_{29}$ = 69%, $N_{7a}$ = 29%, $N_{unassigned}$ = 2%). Temperature: 25°C; Acquisition time: 5 min/FOV; Probes: *hsa*-miR-29_NS_DNA_FP_Cy5_8-nt at 25 nM, *hsa*-miR-16_NS_DNA_FP_Cy5_C9-nt at 75 nM, *hsa*-let-7a_NS_DNA_FP_Cy5_11-nt at 25 nM.

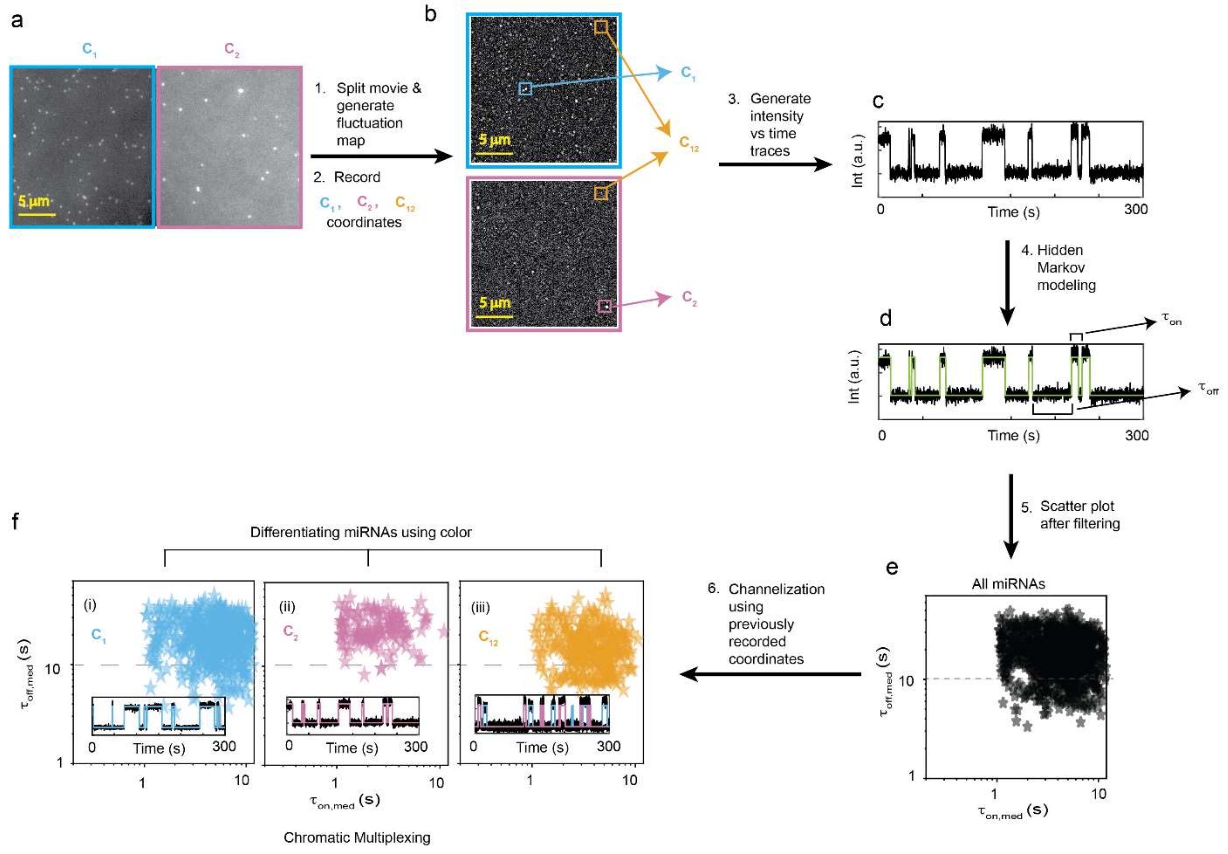


**Supplementary Fig. 2 | Pipeline of Bio-SCOPE with three-color channels. a,** Single-frame images of representative field of view (FOV) from TIRF microscopy. **b,** Intensity fluctuation maps of the fields of view shown in (a). Grey circles indicate the positions of local maxima in the fluctuation map, from which candidate ROIs are identified for further analysis to generate intensity versus time traces. Co-ordinates are recorded for all $C_1$, $C_2$ and $C_{12}$ molecules. **c,** Representative intensity versus time traces generated from the ROIs identified in (b). **d,** Hidden Markov modelling (HMM) idealization (green lines) for each intensity versus time trace shown in (c). Bound and unbound-state dwell times ($\tau_{bound}$ and $\tau_{unbound}$ respectively) are indicated by the horizontal line segments above the idealization. **e,** Scatterplots of dwell time analysis (i.e., $\tau_{on,med}$ versus $\tau_{off,med}$) for all intensity-versus-time trajectories observed within the single field of view. **f,** Demultiplexing of three miRNAs using the previous recorded coordinate information: i) $C_1$ channel, ii) $C_2$ channel and (iii) $C_{12}$ channel. Representative single molecule trace behaviors are shown in the insets. Data from our chromatic multiplexing experiment (shown in Fig 2e) are used here.

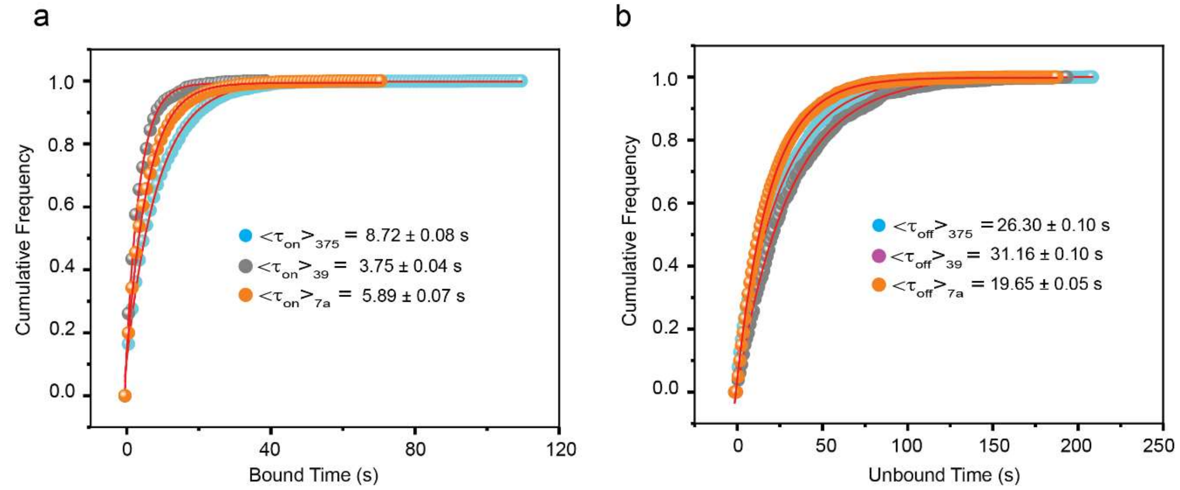


**Supplementary Fig. 3 | Cumulative dwell time analysis of chromatic multiplexing three-plex.** Cumulative (**a**) bound and (**b**) unbound dwell time histograms of *hsa*-miR-375, *cel*-miR-39 and *hsa*-let-7a for chromatic multiplexing three-plex. <$\tau_{on}$> and <$\tau_{off}$> values were determined from single-exponential fitting of cumulative dwell time distributions of accepted single-molecule traces. Temperature: 25°C; Acquisition time: 5 min/FOV; Probes: *hsa*-miR-375_NS_DNA_FP_Cy3_C8-nt at 30 nM, *cel*-miR-39_NS_DNA_FP_Cy5_10-nt at 30 nM, *hsa*-let-7a_NS_DNA_FP_Cy5 & Cy3_11-nt at 30 nM.

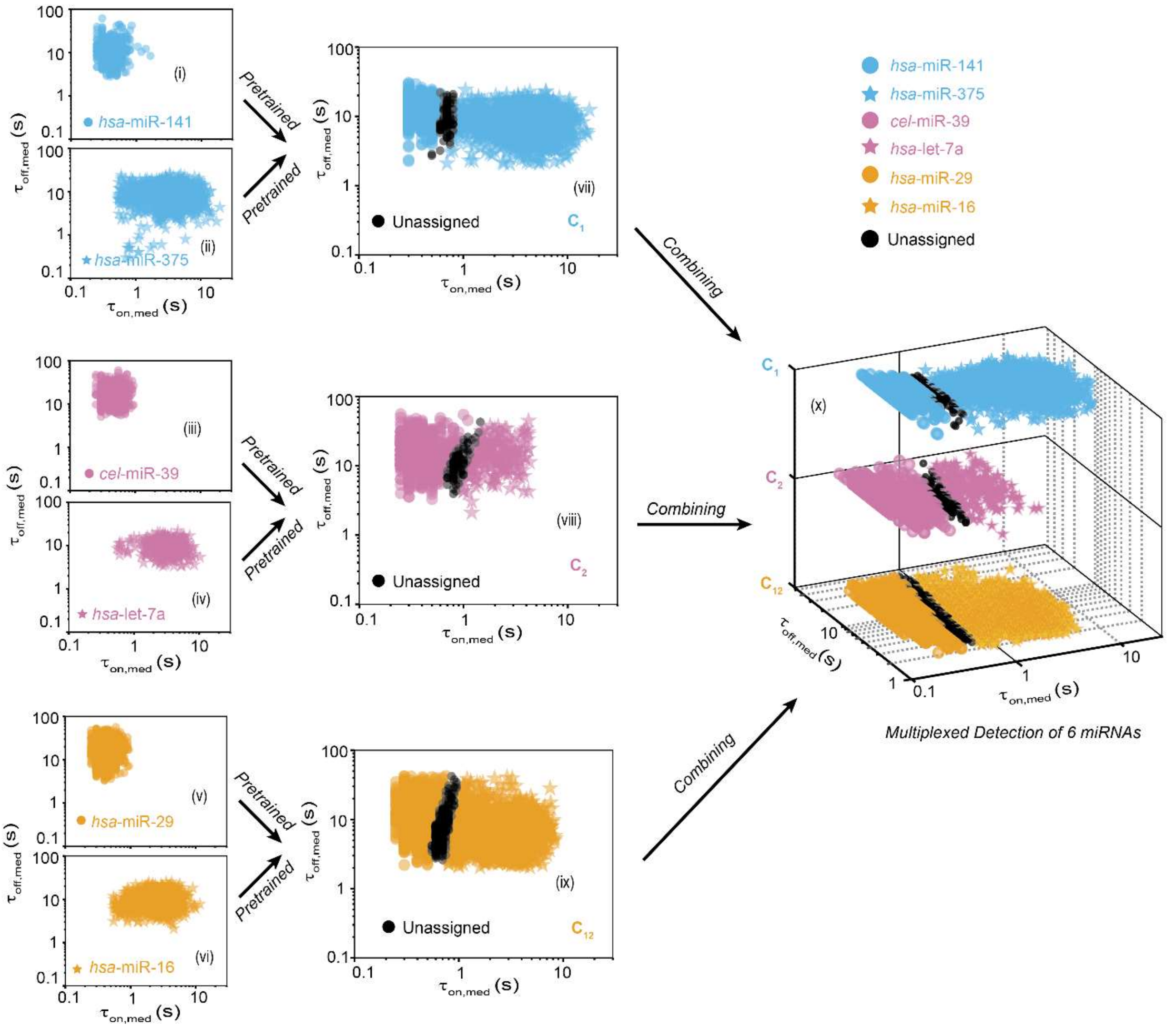


**Supplementary Fig. 4 | Pipeline of Biomarker Single-molecule Chromato-kinetic multi-Omics Profiling and Enumeration (Bio-SCOPE).** Scatterplots of dwell time analysis (i.e., $\tau_{on,med}$ versus $\tau_{off,med}$) for all accepted intensity-versus-time trajectories observed within five fields of view of (i) *hsa*-miR-141, (ii) *hsa*-miR-375, (iii) *cel*-miR-39, (iv) *hsa*-let-7a, (v) *hsa*-miR-29, (vi) *hsa*-miR-16, (vii) a 1:1 mixture of *hsa*-miR-141 and *hsa*-miR-375 ($N_{unassigned}$ =4%), (viii) a 1:1 mixture of *cel*-miR-39 and *hsa*-let-7a ($N_{unassigned}$ =8%), (ix) a 1:1 mixture of *hsa*-miR-29 and *hsa*-miR-16 ($N_{unassigned}$ =7%), (x) multiplexed detection of 6 miRNAs (with 95% confidence clusters). Target concentrations: *hsa*-miR-141: 1 pM, *hsa*-miR-375: 1 pM, *cel*-miR-39: 5 pM, *hsa*-let-7a: 5 pM, *hsa*-miR-29: 1 pM, *hsa*-miR-16: 1 pM; Temperature: 25°C; Acquisition time: 5 min/FOV; Probes: *hsa*-miR-141_NS_DNA_FP_Cy3_8-nt at 25 nM, *hsa*-miR-375_NS_DNA_FP_Cy3_C8-nt at 75 nM, *cel*-miR-39_NS_DNA_FP_Cy5_8-nt at 25 nM, *hsa*-let-7a_NS_DNA_FP_Cy5_11-nt at 75 nM, *hsa*-miR-29_NS_DNA_FP_Cy3 & Cy5_8-nt at 25 nM, *hsa*-miR-16_NS_DNA_FP_Cy3 & Cy5_C10-nt at 75 nM.

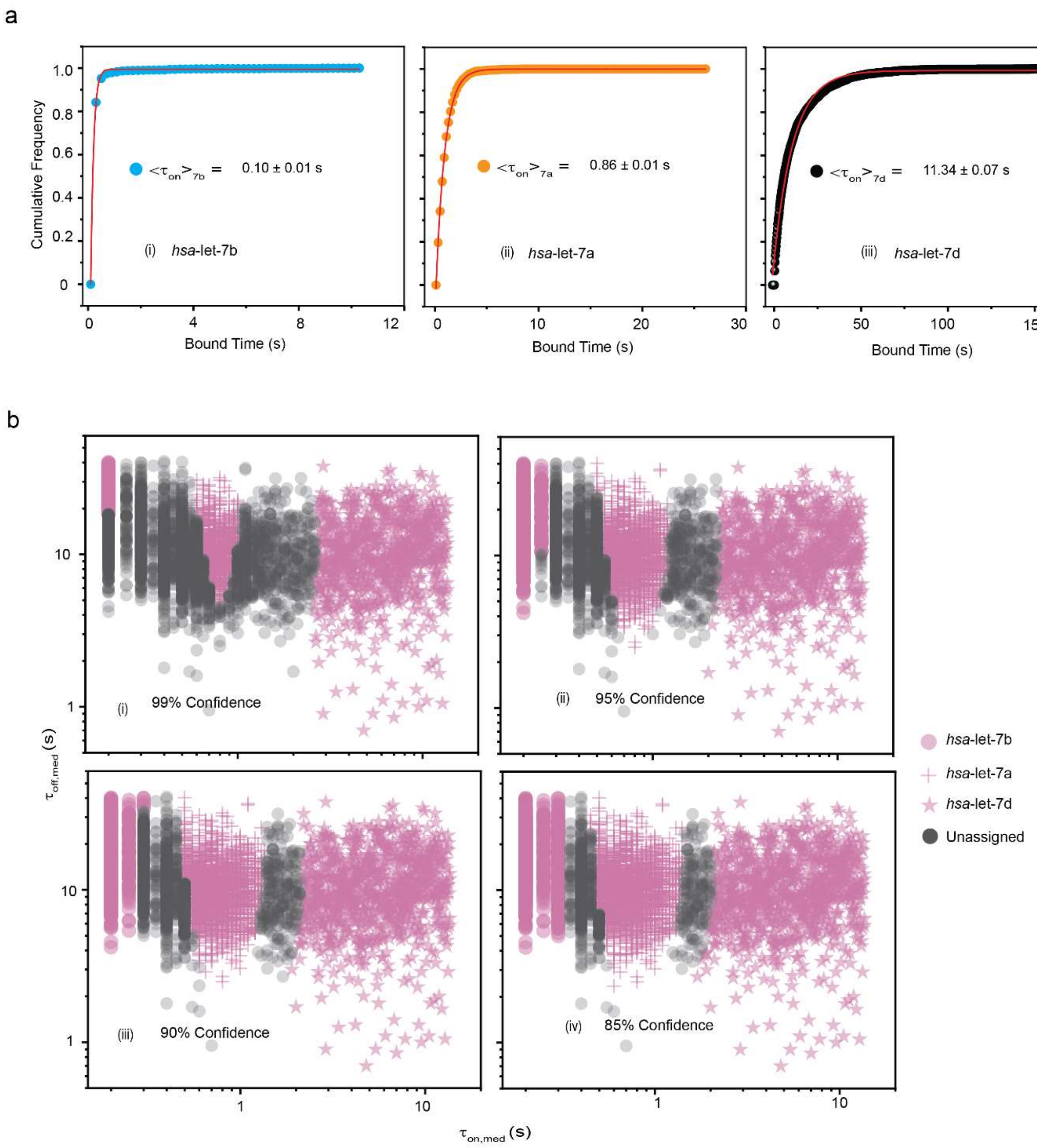


**Supplementary Fig. 5 | Bound times and clustering with different confidence levels of let-7 family three-plex datasets. a,** Cumulative bound time histograms of (i) *hsa*-let-7b, (ii) *hsa*-let-7a, and (iii) *hsa*-let-7d for let-7 family kinetic multiplexing three-plex. $<\tau_{on}>$ values were determined from single-exponential fitting of cumulative bound time distributions of accepted single-molecule traces. **b,** Kinetic multiplexing of the let-7 family three-plex dataset with (i) 99% ($N_{unassigned}$ =56%), (ii) 95% ($N_{unassigned}$ =32%), (iii) 90% ($N_{unassigned}$ =25%) and (iv) 85% ($N_{unassigned}$ =13%) confidence clusters (5 FOVs). Total target concentration: 12 pM (1:1:1); Temperature: 25°C; Acquisition time: 5 min/FOV; Probe: *hsa*-let-7_NS_DNA_FP_Cy5_11-nt at 50 nM.

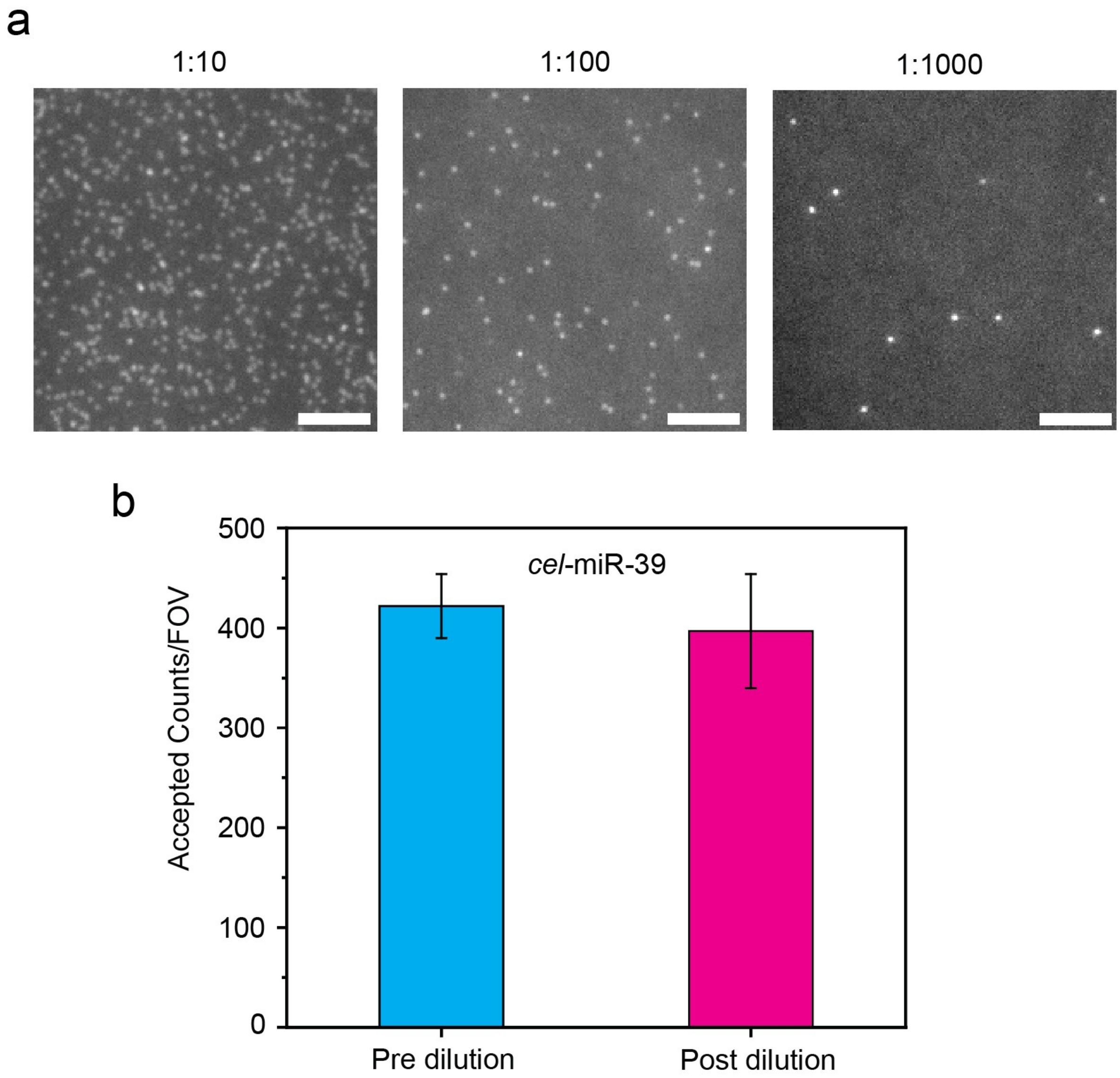


**Supplementary Fig. 6 | Tissue Sample dilution and *cel*-miR-39 doping. a,** Single movie frame of a representative portion of a TIRF microscope field of view (FOV) (scale bar: 5 µm) showing bright puncta at the locations where single fluorescent probes of *hsa*-miR-16 are bound at or near the imaging surface in different dilutions of tissue samples. Sample: human heart tissue total RNA; Probe: *hsa*-miR-16_NS_DNA_FP_Cy5_C10-nt at 50 nM. **b,** Impact of different dilution stages of *cel*-miR-39 doping in tissue samples on accepted counts. Black error bars represent the standard errors of the mean from three independent replicates. Sample: human heart tissue total RNA; Temperature: 25°C; Acquisition time: 5 min/FOV; Probe: *cel*-miR-39_NS_DNA_FP_Cy5_8-nt at 50 nM.

### Supplementary Tables

**Supplementary Table 1 |** miRNA/DNA/Protein sample names and sequences. All sequences are listed 5′-to-3′.

| Sample Type | Name | Sequences |
|---|---|---|
| Targets | *hsa*-miR-141 | 5´/Phos/rUrA rArCrA rCrUrG rUrCrU rGrGrU rArArA rGrArU rGrG/3´ |
| | *hsa*-miR-375 | 5´/Phos/rUrUrUrGrUrUrCrGrUrUrCrGrGrCrUr CrGrCrGrUrGrA/3´ |
| | *cel*-miR-39 | 5´/Phos/rUrC rArCrC rGrGrG rUrGrU rArArA rUrCrA rGrCrU rUrG/3´ |
| | *hsa*-let-7a | 5´/Phos/rUrG rArGrG rUrArG rUrArG rGrUrU rGrUrA rUrArG rUrU/3´ |
| | *hsa*-miR-29 | 5´/Phos/rUrA rGrCrA rCrCrA rUrCrU rGrArA rArUrC rGrGrU rUrA/3´ |
| | *hsa*-miR-16 | 5´/Phos/rUrArGrCrArGrCrArCrGrUrArArArUr ArUrUrGrGrCrG/3´ |
| | *hsa*-miR-21 | 5´/Phos/rUrArGrCrUrUrArUrCrArGrArCrUrGr ArUrGrUrUrGrA/3´ |
| | *hsa*-let-7b | 5´/Phos/rUrG rArGrG rUrArG rUrArG rGrUrU rGrUrG rUrGrG rUrU/3´ |
| | *hsa*-let-7d | 5´/Phos/rArG rArGrG rUrArG rUrArG rGrUrU rGrCrA rUrArG rUrU/3´ |
| | Exon 19 del MUT_22-nt | 5´/TTCCCGTCGCTATCAAGACATC/3´ |
| | T790 MUT_25-nt | 5´/CTCATCATGCAGCTCATGCCCTTCG/3´ |
| LNA Capture Probes (CPs) | *hsa*-miR-141_CP | 5′/A+GAC+A+GT+GTTA /TEG-Biotin/3´ |
| | *hsa*-miR-375_CP | 5′/C+GA +AC+G A+AC +A+AA/TEG-Biotin/3´ |
| | *cel*-miR-39_CP | 5′/A+CC C+GG +TG+A /TEG-Biotin/3´ |
| | *hsa*-let-7_CP | 5' T+AC +T+AC +C+T+C A/3BioTEG/3´ |
| | *hsa*-miR-29_CP | 5´/A+GA+TGGT+GC+TA /TEG-Biotin/3´ |
| | *hsa*-miR-16_non seed CP | 5´/Biotin-TEG/C+GC+CA+AT+AT+TT/3´ |
| | *hsa*-miR-16_seed CP | 5´/C+GT+GC+TGC+TA/TEG-Biotin/3´ |
| | *hsa*-miR-21_CP | 5´C+TG +AT+A A+GC +TA/TEG-Biotin/ 3´ |
| | Exon 19 del MUT_CP | 5´/+AG+CG+ACG+GG+AA/Biotin TEG/3´ |
| | T790 MUT_CP | 5' /Biotin TEG/CG+AAG+GGCAT+G/3' |
| | *hsa* miR 141_ NS_DNA_FP _8-nt | 5'/Cy3/CCATCTTT/3' |
| | *hsa* miR 141_ NS_DNA_FP _10-nt | 5'/Cy3/CCATCTTTAC/3' |
| | *hsa* miR 375_ NS_DNA _FP _8-nt | 5'/Cy3/TCACGCGA/3' |
| | *hsa* miR 375__NS _DNA_FP _C8-nt | 5'/Cy3/CACGCGAG /3' |

| | | |
|---|---|---|
| Fluorescent Probes (FPs) (with Cy5/ Cy3) | *cel*-miR-39_ NS_DNA_FP _8-nt | 5'/Cy5/AAGCTGAT/3' |
| | *cel*-miR-39_ NS_DNA_FP _10-nt | 5'/Cy5/AAGCTGATTT/3' |
| | *hsa*-let-7a_ NS_DNA_FP _10-nt | 5'/Cy5/AACTATACAA/3' |
| | *hsa*-let-7a_ NS_DNA_FP _11-nt | 5'/Cy5/AACTATACAAC/3' |
| | *hsa*-let-7_ NS_DNA_FP _11-nt | 5'/Cy5/AACT ATG CAAC/3' |
| | *hsa*-miR-29_ NS_DNA_FP _8-nt | 5'/Cy5/TA ACC GAT/3' |
| | *hsa*-miR-29_ NS_DNA_FP _9-nt | 5'/Cy5/TAACCGATT/3' |
| | *hsa*-miR-29_ NS_DNA_FP _10-nt | 5'/Cy5/TAACCGATTT/3' |
| | *hsa*-miR-16_ NS_DNA_FP _8-nt | 5'/Cy5/GCCAATAT /3' |
| | *hsa*-miR-16_ NS_DNA_FP _9-nt | 5'/Cy5/GCCAATATT/3' |
| | *hsa*-miR-16_ NS_DNA_FP _C9-nt | 5'/Cy5/CGCCAATAT/3' |
| | *hsa*-miR-16_ NS_DNA_FP _10-nt | 5'/Cy5/GCCAATATTT/3' |
| | *hsa*-miR-16_ NS_DNA_FP _C10-nt | 5'/Cy5/CGCCAATATT/3' |
| | *hsa*-miR-16_S_DNA_FP_8-nt | 5'/TGCTGCTA/Cy3/3' |
| | *hsa*-miR-16_S_RNA_FP_8-nt | 5'/rUrGrCrUrGrCrUrA/Cy3/3' |
| | *hsa*-miR-16_S_RNA_FP_7-nt | 5'/rUrGrCrUrGrCrU/Cy3/3' |
| | *hsa*-miR-21_S_RNA_FP_8-nt | 5'/rArUrArArGrCrUrA/Cy3/3' |
| | Exon 19 del MUT_8-nt | 5'/Cy5/ATGTCTTG/3' |
| | T790 MUT_ G9-nt | 5'/Cy5/GCATGATGA/3' |
| | IL-8_apamer_8A-30 | 5′/Cy5/GGG/i2FU//i2FU/A/i2FU//i2FC/A/i2FU/ /i2FU//i2FC//i2FC/A/i2FU//i2FU//i2FU/ AG/i2FU/G/i2FU//i2FU/A/i2FU/GA/i2FU/AA/3 ' |
| Capture Probe Blockers (CPBs) | *hsa*-miR-141_CPB | 5′/TAACACTGTC/3' |
| | *hsa*-miR-375_CPB | 5′/TTTGTTCGTTC/3' |
| | *cel*-miR-39_CPB | 5'/GGGCCACT/3' |
| | *hsa*-let-7_CPB | 5'/ TGAGGTAGT/3' |
| | *hsa*-miR-29_CPB | 5′/TAGCACCAT C/3' |
| | *hsa*-miR-16_non seed_CPB | 5′/AATATTGGC/3' |
| | *hsa*-miR-16_seed_CPB | 5′/TAGCAGCAC/3' |
| | *hsa*-miR-21_CPB | 5′/TAGCAGCAC/3' |
| | Exon 19 del MUT_CPB | 5'/TTCCCGTCGC/3' |
| | T790 MUT_CPB | 5'/ATGCCCTTCG/3' |
| Inhibitors | *hsa*-miR-16_Inhibitor | 5'/ATCGTCGTGCATTTATAACCGC/3' |
| | *hsa*-miR-21_Inhibitor | 5'/A*C*A*T*C*A*G*T*C*T*G*A*T*A*A*G *C*T/3' ("*" indicates phosphorothioate bonds) |
| Target Blockers (Specific Region) | *hsa*-let-7c_Blocker | 5'/AACCATACAAC/3' |
| | *hsa*-let-7f_Blocker | 5'/AACTATACAAT/3' |
| | *hsa*-let-7g_Blocker | 5'/AACTGTACAAA/3' |
| | *hsa*-let-7i_Blocker | 5'/AACAGCACAAA/3' |
| Carrier | dT 10 | 5'/TTTTTTTTTT/3' |

**Supplementary Table 2 |** Optimized general parameter sets for trace generation and analysis.

| General Trace Generation Parameters | |
|---|---|
| Spot fixing method | Fluctuation map |
| Spot fixing threshold | 3/4 |
| Edge Pixel | 20 |
| Start frame | 1 |
| End frame | 3000 |
| Percentilecut | 0.95 |
| ROI size (pixels) | 3 |

| General Trace Analysis Parameters | |
|---|---|
| Start frame | 1 |
| End frame | 3000 |
| Exposure time (s) | 0.1 |
| Remove_single_frame_events | True |
| Intensity Threshold | 100 |
| S/N Threshold (Event) | 2 |
| S/N Threshold (Trace) | 3 |
| Minimum $N_{b+d}$ | 10 |
| Minimum $N_{b+d}$ | Inf |
| Minimum $\tau_{on,med}$ | 0.2 |
| Maximum $\tau_{on,med}$ | Inf |
| Minimum $\tau_{off,med}$ | 0.2 |
| Maximum $\tau_{off,med}$ | Inf |
| Maximum individual $\tau_{on}$ | Inf |
| Maximum individual $\tau_{off}$ | Inf |
| Maximum $\tau_{on}$ (C.V.) | Inf |
| Maximum $\tau_{off}$ (C.V.) | Inf |
